# Dissecting the regulatory activity and sequence content of loci with exceptional numbers of transcription factor associations

**DOI:** 10.1101/2019.12.21.885830

**Authors:** Ryne C. Ramaker, Andrew A. Hardigan, Say-Tar Goh, E. Christopher Partridge, Barbara Wold, Sara J. Cooper, Richard M. Myers

**Affiliations:** HudsonAlpha Institute for Biotechnology, Huntsville, AL; University of Alabama at Birmingham, Department of Genetics, Birmingham, AL; California Institute of Technology, Division of Biology and Biological Engineering, Pasadena, CA

## Abstract

DNA associated proteins (DAPs) classically regulate gene expression by binding to regulatory loci such as enhancers or promoters. As expanding catalogs of genome-wide DAP binding maps reveal thousands of loci that, unlike the majority of conventional enhancers and promoters, associate with dozens of different DAPs with apparently little regard for motif preference, an understanding of DAP association and coordination at such regulatory loci is essential to deciphering how these regions contribute to normal development and disease. In this study, we aggregated publicly available ChIP-seq data from 469 human DAPs assayed in three cell lines and integrated these data with an orthogonal dataset of 352 non-redundant, *in vitro*-derived motifs mapped to the genome within DNase hypersensitivity footprints in an effort to characterize regions of the genome that have exceptionally high numbers of DAP associations. We subsequently performed a massively parallel mutagenesis assay to search for sequence elements driving transcriptional activity at such loci and explored plausible biological mechanisms underlying their formation. We establish a generalizable definition for High Occupancy Target (HOT) loci and identify putative driver DAP motifs in HEPG2 cells, including HNF4A, SP1, SP5, and ETV4, that are highly prevalent and exhibit sequence conservation at HOT loci. The number of different DAPs associated with an element is positively associated with evidence of regulatory activity and, by systematically mutating 245 HOT loci, we localized regulatory activity to a central core region that depends on the motif sequences of our previously nominated driver DAPs. In sum, this work leverages the increasingly large number of DAP motif and ChIP-seq data publicly available to explore how DAP associations contribute to genome-wide transcriptional regulation.

## Introduction

Gene expression networks underlie many cellular processes (Spitz and Furlong 2012). These expression networks are controlled in cis by DNA regulatory elements, such as promoters and enhancers, which can be proximal, distal, or within their target genes in a given expression network. Extensive mapping of epigenetic modifications and 3D chromatin structure have provided an increasingly rich set of clues to the locations and physical connections among such elements. Nevertheless, these biochemical signatures cannot yet accurately predict the presence or amount of regulatory activity encoded in underlying DNA. There are many known and suspected reasons for this difficulty, including the relative strength, number of interacting partners, and redundancy of each element, each of which may modulate a locus’ contribution to the native expression level(s) of its respective target gene(s) in a manner difficult to predict without direct experimentation (Roadmap Epigenomics Consortium 2015; The ENCODE Project Consortium 2007, 2012; Sanyal et al. 2012). In this manuscript, we present evidence that the total number of DNA-associated proteins (DAPs) that associate with a locus can act as a quantitative predictor of the locus’ regulatory activity and that the activities of loci with large numbers of DAP associations can be disrupted in a predictable manner by altering subsets of putative “driver motifs”.

Classically, regulatory loci are thought to be discriminately bound by a small subset of expressed transcription factors (i.e. <10) in a manner governed by each factor’s DNA sequence preference, and additional proteins are recruited through specific protein-protein interactions (Mitchell and Tjian 1989). However, this model is becoming incongruent with observed DAP associations as catalogs of genome-wide DAP binding maps continue to expand (Foley and Sidow 2013). Specifically, the discriminatory nature by which regulatory regions recruit DAPs is unclear at thousands of loci that have been shown to associate with dozens of different DAPs with seemingly no regard for motif preferences (Ramaker et al. 2017; Wreczycka et al. 2019; Teytelman et al. 2013; Jain et al. 2015). These loci, which have associations with dozens of DAPs, have been inconsistently defined, but are broadly referred to as high occupancy target (HOT) sites. This phenomenon has been at least partly attributed to technical artifacts of chromatin immunoprecipitation sequencing (ChIP-seq), a common assay used to map DNA-protein interactions *in vivo*, resulting in a small number of blacklisted loci (Johnson et al. 2007; Landt and Marinov 2012; Carroll et al. 2014). These artifacts have largely been localized to regions of the genome to which it is difficult to confidently align sequencing reads, such as repetitive elements (Carroll et al. 2014; Landt and Marinov 2012). Others found potentially misleading or non-specific ChIP-seq signal more broadly at GC-rich promoters of highly expressed genes (Wreczycka et al. 2019). These findings provide motivation for proceeding with caution when analyzing DAP co-associations, particularly at HOT loci. In our analysis of these genomic regions, we present an extensive examination of potentially confounding characteristics of HOT loci, employed conservative peak calling thresholds standardized by the ENCODE consortium, and rely heavily on orthogonal, non ChIP-seq based, data sets to define DAP associations. Despite this conservative approach, we find complex DAP co-associations to be pervasive throughout the genome, and the increasing completeness of our catalog of DAP occupancy maps, generated by ChIP-seq and other orthogonal approaches, invites a systematic investigation of the prevalence and significance of DAP co-associations and of the classic model for how DAPs interact with regulatory elements.

Previous work has investigated HOT loci using a combination of genome-wide transcription factor motif scanning and ChIP-seq experiments (Foley and Sidow 2013; Li et al. 2015, 2016). These studies have found thousands of loci harboring dozens of motif or ChIP-seq peak-based, DAP associations throughout the genome and have labeled these regions as High Occupancy Target (HOT) loci. HOT loci were found to be enriched for markers of regulatory activity, such as initiating *POLR2* binding, DNase hypersensitivity, active histone marks, and strong activity in enhancer reporter assays conducted in transgenic mouse embryos. These studies also demonstrated that context-specific HOT loci are generated in association with cell differentiation and oncogenesis at locations enriched for disease-risk variants. However, these studies were largely limited to experimental data from fewer than 100 transcription factors derived from several different cell lines. Previous studies also have not incorporated genome-wide 3D chromatin structure data, such as Chromatin Interaction Analysis by Paired-End Tag sequencing (ChIA-PET) and promoter capture Hi-C experiments, which have been performed on an increasing number of cell lines. Massively parallel reporter assay (MPRA) data that probes the regulatory activity at thousands of loci across the genome is also now available to assess for quantitative correlation with DAP associations. Furthermore, few studies have performed experimental perturbations on HOT loci to quantify their vulnerability to single base pair mutations and to probe the key sequence features driving their activity.

In this manuscript, we aggregated ChIP-seq data from 469 DAPs assayed in three cell lines. We integrate these *in vivo* mapping data with an orthogonal dataset of 352 non-redundant, *in vitro*-derived motifs from 555 DAPs, which we have mapped to the genome within DNase hypersensitivity footprints for each cell line. Specifically, we aim to (1) detail the prevalence and cell type-specificity of regulatory element DAP co-associations, (2) assess the utility of co-associations as a marker of regulatory activity, (3) perform a high-resolution dissection of key sequences driving activity at regions with large numbers of DAPs by using a massively parallel mutagenesis assay, and (4) explore potential factors influencing observed DAP co-associations, such as 3D chromatin interactions, sequence content and copy number variation.

## Results

### HOT loci are prevalent in the genome

We used two orthogonal methods to infer DAP associations across the genome. The first involved analysis of ENCODE ChIP-seq peaks (208, 129, 312 DAPs in the HepG2, GM12878, K562 cell lines; Table S1). A subset of DAPs was further classified into sequence specific transcription factors (ssTFs, *N*=117) and non-sequence specific DAPs (nssDAPs, *N*=85). ssTFs were conservatively defined as those that had an *in vitro* derived motif in the CIS-BP database (Weirauch et al. 2014) and nssDAPs were defined as DAPs without *in vitro* derived motifs that had previously been characterized as non-sequence specific chromatin regulators or transcription cofactors (Partridge, 2018, Lambert, 2018). As a second method to assess transcription factor associations, we used the Protein Interaction Quantitation (PIQ) algorithm and *in vitro* derived (SELEX, protein binding microarray, or B1H) motifs from 555 TFs in the CIS-BP database to identify DAP footprints that were present in ENCODE DNase I hypersensitivity (DHS) footprints (Table S2) (Sherwood et al. 2014). To quantify DAP co-associations, we binned the genome into a minimal set of non-overlapping 2-kb loci that encompassed either every ChIP-seq peak or every distinct DHS footprint and counted the number of unique DAP peaks or footprinted motifs contained within each locus (Table S3-6). We focused on HepG2 as the primary cell line in our analysis and the figures in this paper contain HepG2-derived data unless otherwise specified.

To ensure that our definition of a “HOT” locus was generalizable across cell lines and datasets, we defined HOT regions as those associated with at least 25% of DAPs assayed. This definition requires 52 of 208 DAPs assayed with ChIP-seq in the HepG2 cell line to have a peak at a given locus to reach the HOT threshold. Nearly 6% of loci (13,792 out of 244,904) met this HOT threshold in HepG2, and we found this result to be consistent after varying the number of DAPs incorporated into our analysis via random sampling (Figure 1A, Figure S1A-C). We found our 25% threshold to be preferable to other HOT thresholds based on the stability of number of loci detected and recall performance of the full data set in a series of random down-samples (Figure S1B-C). This threshold also performed similarly in the K562 and GM12878 cell lines (Figure S1D-F). The distribution of observed DAP co-associations was dramatically different than that observed after randomly scrambling DAPs across all loci (K-S test P< 5E-16), with no locus reaching our HOT threshold by random chance (Figure S1G). A subset of our HOT loci fall under a previously established definition of super enhancers (Whyte et al. 2013); however, no HOT promoters and the vast majority (97.1%) of HOT enhancers are not encompassed by this definition (Table S7, Figure S1H-I). The observed pattern of DAP co-associations was relatively consistent when restricting to ssTFs or nssDAPs (Figure 1B, S1A), although a slightly larger proportion of ssTF peaks were found at a locus alone (44.0% vs. 34.4%) or with a relatively small number of co-associated ssTFs. No locus had 25% or more of the 352 non-redundant HepG2 <u>D</u>HS-<u>F</u>ootprinted <u>M</u>otifs (DFMs) analyzed, suggesting the number of possible motifs at a locus is constrained in a manner not observed for ChIP-seq peaks. However, the number of DFMs was positively correlated with the number of gross DAP peaks across loci (rho=0.494, P<5E-16) despite a minority (∼10%) of ssTFs with ChIP-seq peaks present at any given HOT site possessing a corresponding DFM at the same loci (Figure 1C). These data suggest that, although the presence of DFMs is a strong indicator of HOT loci, a majority of DAP associations at HOT loci likely represent non-specific or indirect interactions.

**Figure 1.**
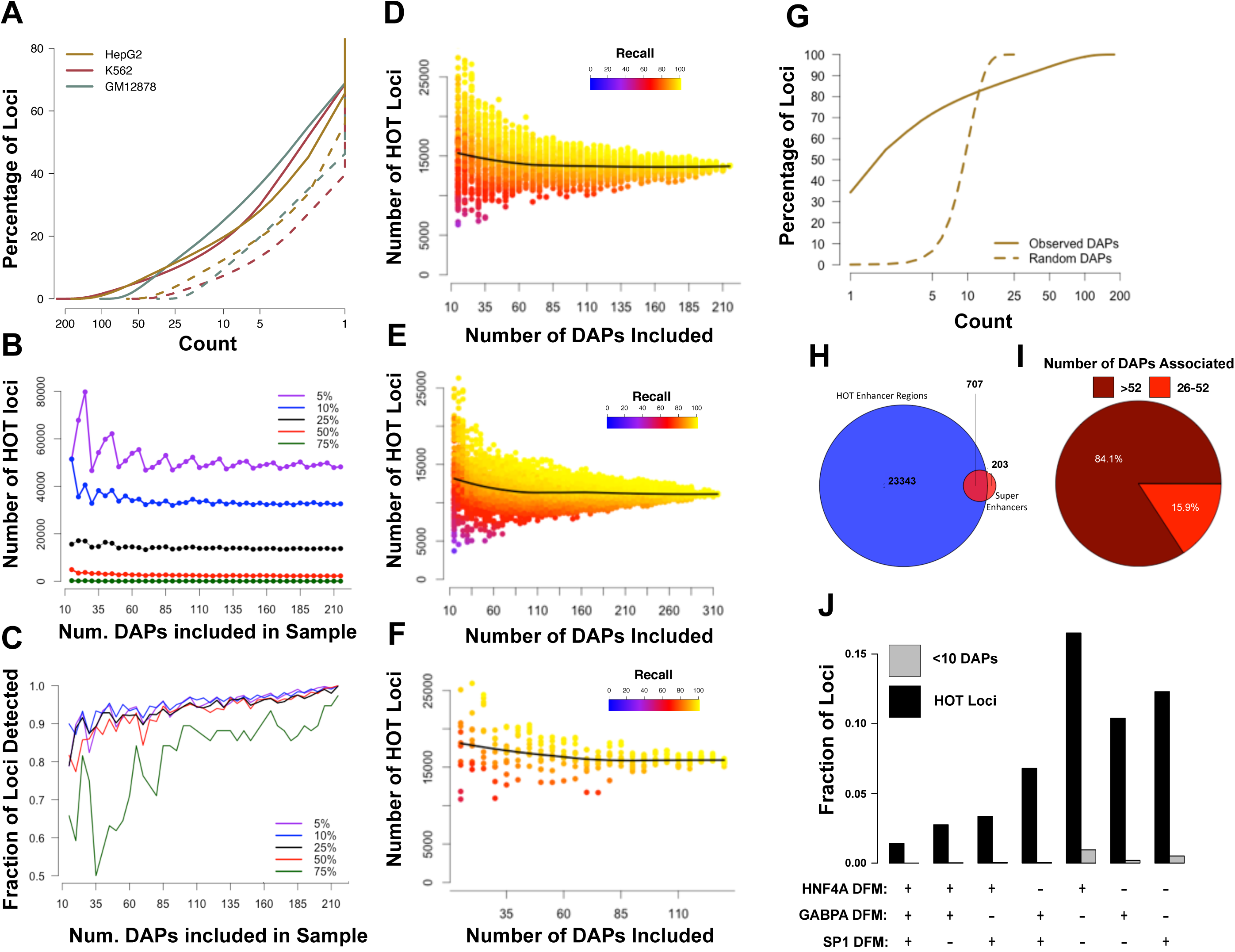
(A) Number of loci reaching “HOT” threshold of 25% of unique ChIP-seq peaks after performing random down sampling (from the original 208) of the number of DAPs included. Each data point represents the result of a random sampling of a specified number DAPs. The color indicates the recall performance or the percentage of true HOT sites, as defined by >25% of DAPs bound in the full dataset, detected with current sample of DAPs. The black line represents the median result of 100 random samples of each number of DAPs as specified by the x-axis. (B) Cumulative distribution function (CDF) showing the proportion of loci containing at least a given number of unique DAP ChIP-seq peaks in HepG2. The green line shows data for all 208 DAPs, the red dashed line shows data for nssDAPs, and the yellow dashed line shows data for ssTFs. (C) Boxplots demonstrating the number of ChIP-defined DAPs with a corresponding DFM present at the same locus at various levels of DAP co-association. (D) Barplots indicating the fraction of ChIP peaks for each DAP that fall within HOT loci. Bars are grouped by previously defined DAP classes. The dashed red line indicates the average fraction (55%) of ChIP peaks that fall within a HOT locus across all DAPs. (E) Scatter plot demonstrating the fraction of HOT sites that contain a ssTF ChIP-seq peak and a DFM. ssTFs highlighted in the top right are putative driver TFs present at high proportion of HOT sites.

While HOT sites represent a small minority of DAP-associated loci, due to the massive number of DAPs that localize to these sites, they account for 55% of any individual DAP’s ChIP-seq peaks on average, potentially complicating the interpretation of any individual ChIP-seq dataset. We observed a wide range in the rate of participation in HOT loci within previously defined DAP classes, but DAPs with a methyl binding domain (MBD), a Myb/SANT domain, or a homeodomain exhibit the highest rates of HOT site participation (Figure 1D). These classes have been previously described as having an affinity for large multi-protein complex membership, such as the NuRD complex, and are plausible candidates to be indirectly recruited to HOT loci (Basta and Rauchman 2009; Underhill et al. 2000). A small number of ssTFs, including SP1 and SP5, which bind GC rich sequences, HNF4A, a key driver of liver cell differentiation, GABPA and ETV4, which belong to the ETS family of ssTFs, and KLF16 had DFMs at an exceptional number of HOT sites (Figure 1E) (Wei et al. 2010; Tan and Khachigian 2009; DeLaForest et al. 2011). Many of these ssTFs have been implicated as drivers of liver expression programs, and thus can be reasonably nominated as putative “drivers” of HOT sites in HepG2, a liver cancer-derived cell line (DeLaForest et al. 2011). Despite rampant co-associations of DFMs, we observed little evidence for specific cooperation among these driver ssTFs as HOT loci were roughly three times more likely to have only one of HNF4A, GABPA or SP1 DFMs present rather than any combination of the three (Figure S1J).

### HOT loci are enriched for promoter and enhancer regions near highly expressed genes

After establishing the prevalence of HOT loci, we investigated the biological significance of loci with a large number of DAPs. Intersecting these loci with previously assigned HepG2 genomic annotations, we found a continuous relationship between the number of DAPs, identified as ChIP-seq peaks or DFMs, and enhancer or promoter designation from the IDEAS genome segmentation algorithm (Figure 2A, S2A) (Zhang et al. 2016). Loci containing a large number of DFMs were particularly enriched for promoters over other annotations (Figure S2A). Roughly half of all IDEAS promoters and likely enhancers in HepG2 met our HOT loci threshold, while genomic regions with other annotations rarely met this threshold (Figure S2B). Less than 2% of loci without an enhancer or promoter annotation met our HOT threshold (Figure S2B).

**Figure 2.**
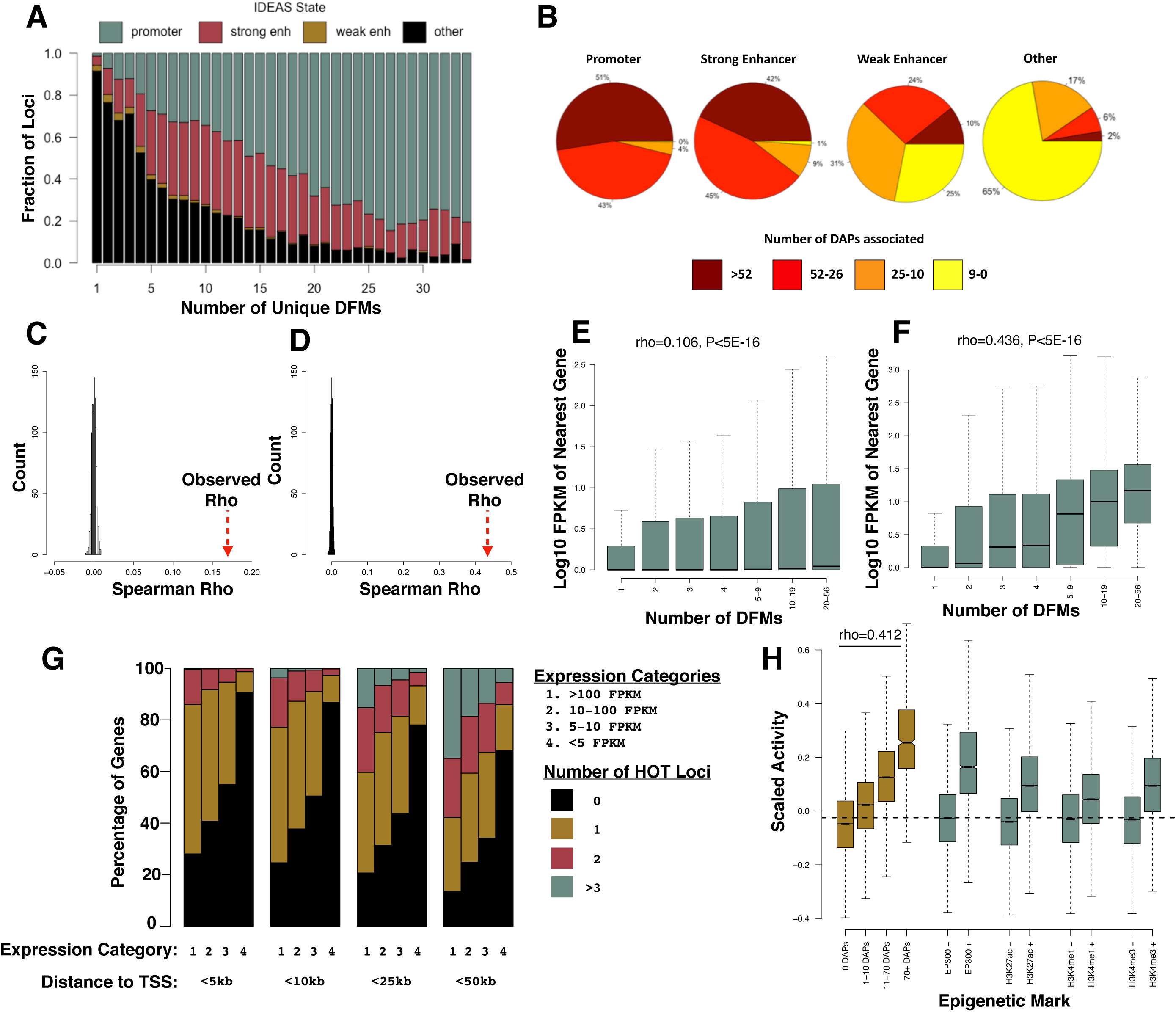
(A) IDEAS annotations of loci binned by ChIP-defined DAP associations. Promoter, strong enhancer, and weak enhancer annotations represent 0.27%, 0.35%, and 0.22% of the HepG2 genome, while the remaining 99.16% of the genome (largely consisting of quiescent and repressed annotations) was used for the “Other” annotation. (B-C) The expression level of the maximally expressed gene neighboring each locus binned by the number of ChIP-defined DAP associations. Plots show loci either distal (>5kb, B) or proximal (<5kb,C) to their nearest gene. The sample size of each bin is as follows: 1-9 (*N*=194028), 10-19 (*N*=17148), 20-29 (*N*=8685), 30-39 (*N*=5876), 40-49 (*N*=4578), 50-69 (*N*=6532), 70-99 (*N*=5351), 100+ (*N*=2706). (D) ChIP- and DFM-defined co-association correlates with activity in a previous high throughput reporter assay conducted on ∼2000 selected enhancer regions in HepG2. (E) Scatter plots demonstrating the fraction of distal (>5kb from a TSS) and proximal (<5kb from a TSS) HOT sites that contain a DFM for each ssTF in HepG2.

To assess the regulatory activity of loci as a function of the number of unique DAPs, we used a variety of publicly available gene expression and reporter activity datasets. Using ENCODE HepG2 RNA-sequencing data, we found a positive association between the number of unique DAPs at a locus and the maximum expression level of nearby genes, particularly in loci proximal (<5kb, rho=0.436, P<5E-16) to a transcription start site (Figure 2B-C, S2C-F). Specifically, 55% of genes whose TSS were less than 5kb from a HOT locus were expressed at a level of 10 FPKM or higher, while only 16% of genes near a locus with fewer than 10 DAPs bound exhibited a similar expression level. Highly expressed genes, with FPKMs greater than 100, were also three times more likely to have multiple HOT loci within 50kb of their TSS than genes with FPKMs less than five (chi-square, P<5E-16, Figure S2G). Loci distal to a TSS exhibited a significantly weaker correlation (Fisher r-to-Z transformation = 67.87, P<5E-16, Figure 2B). Both ChIP-seq and DHS motif-defined (Figure 2D) DAP associations positively correlated with activity in previous high-throughput reporter assays of ∼2000 selected loci in HepG2 and in ATAC-seq fragments in GM12878 (rho=0.230 and 0.207, P<5E-16, Figure S2H) (Inoue et al. 2017; Wang et al. 2018). For both reporter assay datasets, the number of DAPs represents a specific, quantitative marker of regulatory activity that compares favorably to commonly used markers of promoter or enhancer activity (Figure 2D, S2H).

A small number of DFMs exhibited a preference for loci distal (>5kb) or proximal (<5kb) to a TSS (Figure 2E). Specifically, HNF4A, NR2F6, JDP2, and FOX family motifs demonstrated a 2-fold preference for distal, enhancer HOT loci and ETS and SP family motifs had a 3-fold bias for proximal, promoter HOT loci. These findings agree with previous studies that have found HNF4A occupancy at enhancers to be essential for activity in mouse hepatocytes (Thakur et al. 2019) and a strong promoter bias for the ETS family of motifs (Hollenhorst et al. 2007). The level of sequence conservation of driver TF motifs was higher in HOT loci (Figure S3A) and the degree of both TSS-distal and TSS-proximal motif conservation was correlated with total number of DAPs at a locus (Figure S3B-C). This correlation was not observed for CTCF motif (Figure S3B-C). In sum, these data suggest a dose-dependent relationship between the number of DAPs and the regulatory activity of a locus. This relationship is relatively unchanged after restricting analyses to ssTFs or nssDAPs, although nssDAPs tended to be slightly more predictive of activity than did ssTFs (Table S8).

### High-throughput mutagenesis of HOT loci reveals motifs driving activity and possible mutational buffering

After establishing that HOT loci exhibit strong regulatory activity in a variety of reporter assays, we next sought to explore the key sequence features driving this activity by performing experimental perturbations of the sequence content of several loci. A naive hypothesis for how sequence motifs contribute to activity at HOT loci is an additive one, in which the regulatory activity of a locus is simply the sum of each constituent motif’s contribution. In this scenario, ablation of a motif would have a roughly equivalent effect across loci regardless of neighboring sequence content. A more sophisticated model allows for interactions between constituent motifs with synergistic or redundant relationships. If motif synergy is a prominent feature, one would expect individual motif disruptions to have a greater effect on activity in loci containing large numbers of motifs while the opposite would be true if motif redundancy was the predominant relationship between motifs. Alternatively, it is possible regulatory activity at HOT loci is not wholly dependent upon individual motifs and is substantially derived from other features. In that situation the most disruptive mutations would not map to known TF motifs.

To begin to resolve these competing models and to identify the sequence elements most important in controlling regulatory activity at HOT loci, we performed a Self-Transcribing Active Regulatory Region Sequencing (STARR-seq)-based mutagenesis assay on 245 genomic loci that had previously demonstrated activity in massively parallel or single locus reporter assays (Table S9). Assayed loci contained a range of unique ChIP-seq peaks (1-150 unique DAP peaks), although the vast majority met our HOT loci threshold by containing called peaks for 52 or more DAPs. Within each 2-kb locus, we designed oligos centered around a 390-bp region of maximal ChIP-seq signal intensity across all DAP peaks (Figure 3A). We found a majority of ChIP-seq peaks and DFMs localized to a few hundred base pairs within each HOT loci bin (Figure S4A), and thus we reasoned this approach would capture a majority of active elements within each locus and allow us to assay several different loci. Each 130-bp oligo represented a left, right, or central window of the 390-bp core region. For the positive strand, we synthesized reference sequence for each window in addition to tiled 5-bp (AAAAA or TTTTT, depending on maximal disruption from reference sequence) mutations. For the central 130-bp window, we also included oligonucleotides with tiled single base pair mutations at each position in addition to the tiled 5-bp mutations for both the positive and reverse strand. Control sequences consisting of oligonucleotides matched for GC content and repeat length and previously tested null sequences were also included in our library.

**Figure 3.**
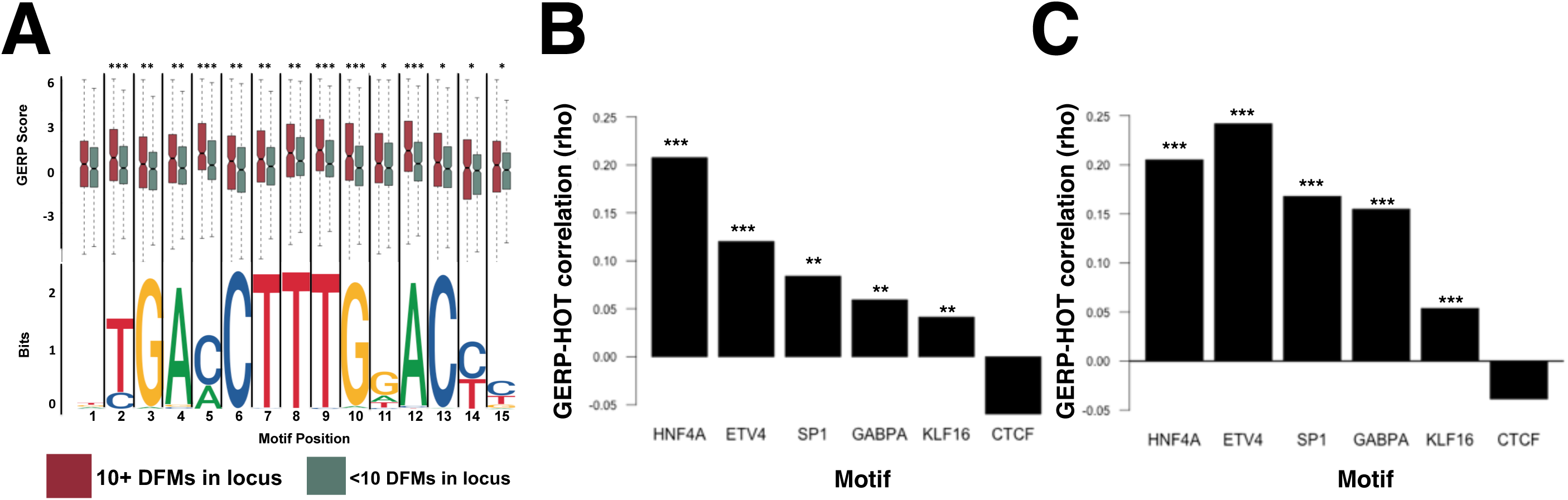
(A) Example locus depicting mutagenesis schema. The red region indicates a 130bp core, centered upon the maximum number of unique ChIP-seq peaks and DFMs, in which we performed tiled single bp and five bp mutagenesis in both the forward and reverse orientation. The flanking green regions represent 130bp sequences flanking the core region in which we performed tiled 5bp mutagenesis in the forward orientation only. (B) Boxplots indicating activity (as represented by the RNA/DNA ratio) was largely concentrated in the WT core loci in both the forward and reverse orientations and not in flanking regions or null regions (Wilcoxon P<5E-16). (C) Plot indicating the proportion of mutations imposing a change of activity at a variety of thresholds for five bp mutations. Green points indicate data for mutations falling within DHS footprints. Red points indicate data for mutations falling outside of DHS footprints. (*) Indicates Fisher’s P<0.05. (D) Scatter plot demonstrating the fraction of HOT sites that contain an ssTF ChIP peak and DFM. TFs highlighted in the top right are putative direct binding TFs associated with a high proportion of HOT sites. The size of each point corresponds to an ssTFs DFM enrichment for high impact mutations in our mutagenesis assay. (E) Barplot showing the cumulative differential activity (locus mean – mutation) across all positions in the HNF4A motif. (F) Scatterplot demonstrating the number of non-redundant DFMs at a locus is inversely correlated with its vulnerability to mutation (expressed as the sum of all mutation delta activity scores).

We cloned oligonucleotides into the STARR-seq reporter vector and transfected the plasmids into HepG2 cells. We subsequently collected RNA from transfected cells to assess the relative abundance (and thus activity) of each test element compared to DNA library input. We detected more than 90% of individual elements post-transfection (Figure S4B-C) and observed that poorly represented elements were evenly distributed in position across each locus, and thus were likely not a product of alignment efficiency (Figure S4D). With the exception of a subset of mutated sequences, RNA and DNA counts were highly correlated across our element library (rho=0.955, P<5E-16, Figure S4E-H). RNA/DNA ratios were also highly correlated across sequencing replicates at our conservative minimum representation threshold of two DNA counts per million (CPM) (Figure S4I-J). As expected, elements from the central window were significantly more active (higher RNA/DNA ratio) than those on the border of regions of ChIP-seq signal (Figure 3B, S4E). Elements with single base pair mutations exhibited roughly equivalent activity to those with reference sequence on average, but displayed a greater range in activity (Figure S5A). This suggests that, except for a small subset, most single base mutations did not significantly affect activity. Elements with 5-bp mutations exhibited slightly less activity than reference sequence elements on average (Wilcoxon P<5E-16, Figure S5A). We found the effects on activity of most mutations were highly correlated between strands (rho=0.45, P<5E-16, Figure S5B-C) and transversions tended to have more impact than transitions, as previously reported (Figure S5D) (Guo et al. 2017). Furthermore, we successfully validated 14 high-impact mutations (including one gain-of-activity mutation) and 14 adjacent low-impact control mutations with individual luciferase reporter experiments using two different plasmids that place the test element either upstream or downstream of the reporter (Figure S6-S7, Table S10).

Mutations that affect previously defined DFMs showed the greatest effect on test element activity (Figure 3C, S5E) and the magnitude of mutation effects was strongly correlated with that predicted by LS-GKM, an algorithm developed for predicting mutation effects on TF motifs (Lee 2016) (rho=0.304, P<5E-16, Figure S5F). Thus, activity at loci with large numbers of DAPs associated seem to be controlled by conventional recognition motifs that can be disrupted in a predictable manner. A motif’s predilection for impactful mutations was associated, but weakly, with its overall enrichment at HOT loci (rho = 0.320, P=0.022, Figure 3D). Of particular interest are ETV4, SP1, SP5, and HNF4A, each of which is highly prevalent across all HOT loci and enriched for high impact SNVs, providing further evidence that these ssTFs may be important drivers of activity at HOT loci (Figure 3D). Broadening our enrichment analysis to include all CIS-BP motifs of ssTFs expressed in HepG2, not just those assayed by ChIP-seq, reveals several additional ssTF motifs strongly enriched for high-impact mutations such as the AP-1 and FOXA sequences (Table S11). The resolution of our mutagenesis assay allows us to identify the most important base pairs governing activity in each of these motif sequences (Figure 3E, Figure S5G-H). We also found evidence of partial motif redundancy, as loci with high numbers of motifs were generally less vulnerable to single nucleotide variation (rho=-0.271,P=4.1E-5, Figure 3F). This suggests that some HOT loci are potentially buffered from motif disrupting mutations that could completely ablate other loci with fewer motifs. Independent support for this hypothesis comes from the observation that the effect sizes of significant eQTL SNPs mapping to HOT loci tend to be significantly lower (rho=-0.175, P<5E-16, Figure S8) (Varshney et al. 2019).

### HOT loci dichotomize into cell-type specific or ubiquitous groups

Integrating data from multiple cell lines allowed us to examine the cell type-specificity of HOT loci and corresponding DAP associations. Loci containing an increasing number of unique ChIP-seq peaks in HepG2 were more likely to be present in both K562 and GM12878 than were loci with fewer ChIP-seq peaks (Figure S9A-B). HOT sites across each cell line tended to fall within two groups: one in which HOT sites were present in only one cell line, and a smaller group in which sites were present in all three cell lines (Figure 4A). Relatively few loci were present in only two of three cell lines.

**Figure 4.**
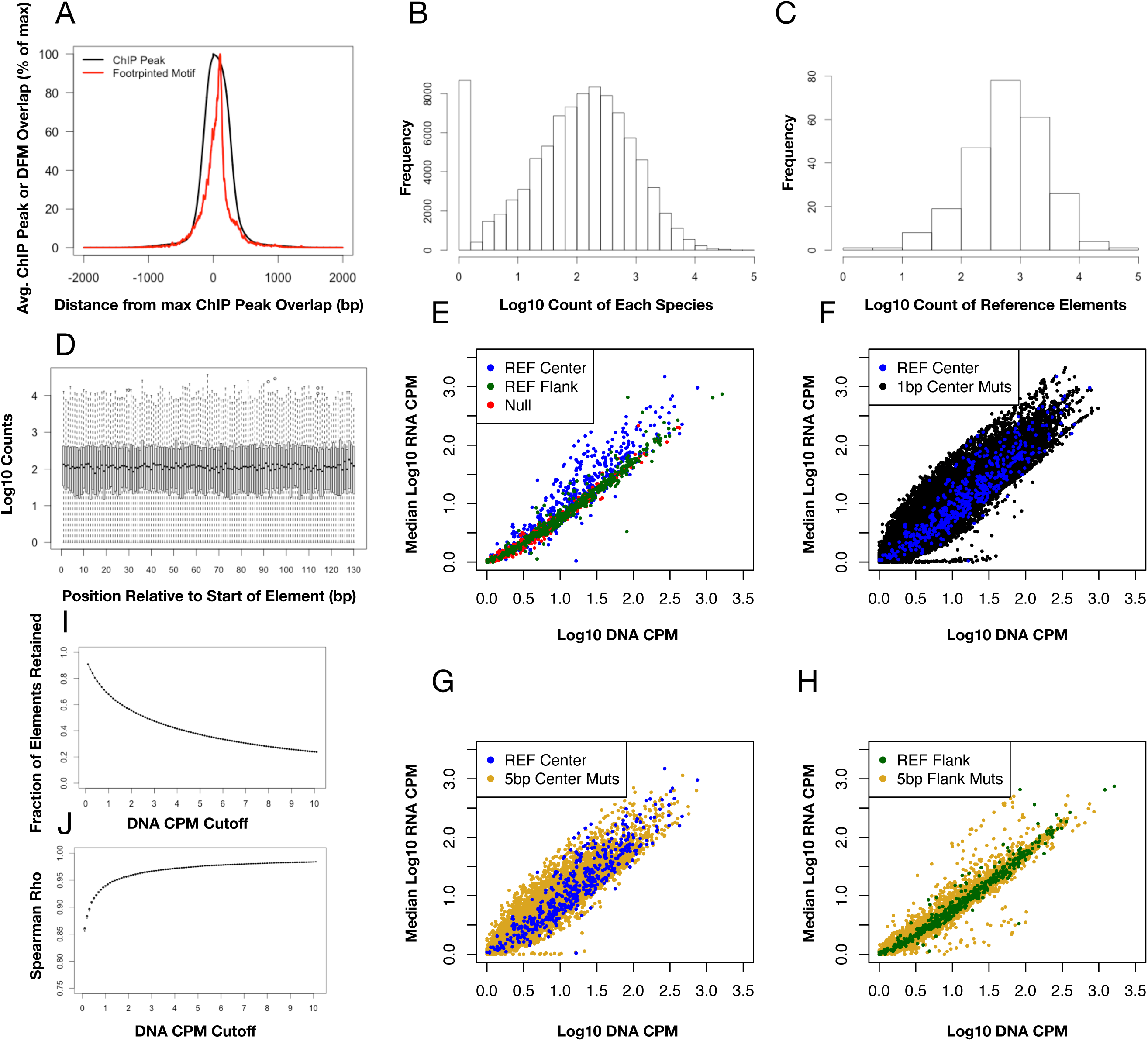
(A) The number of HOT loci present in all possible combinations of each cell line. (B) Scatter plot demonstrating the association between cell type-specific, HOT loci enrichment and distal, HOT loci enrichment in HepG2. Cell type-specificity enrichment value is computed by subtracting the fraction of HepG2-specific HOT loci (N=7692) in which a DFM is present from the fraction of non HepG2-specific HOT loci (N=6100) in which a DFM is present. The distal locus enrichment is computed by subtracting the fraction of HOT loci >5kb from the nearest TSS (N=6445) in which a DFM is present from the fraction of HOT loci <5kb from the nearest TSS in which a DFM is present (N=7347). (C) Stacked bar plots displaying the proportion of cell type-specific or expression level matched, non-cell type-specific genes that possess a specified number of neighboring cell type-specific or non-cell type-specific HOT loci at a specified distance to TSS threshold. Cell type-specific genes were computed by randomly sampling 500 genes that were expressed at least 4 fold higher in the cell line of interest than the other two cell lines and had an FPKM of at least five in the cell line of interest. Non-cell type-specific genes were a cell type-specific gene expression level matched sample of 500 genes with an FPKM of at least five in HepG2, K562, and GM12878. (D-E) Proposed model of how HOT loci relate to cell type-specific (D) and non-cell type-specific housekeeper gene (E) expression.

We found DFMs that were biased towards distal HOT loci in Figure 2E (HNF4A, NR2F6, and FOXA family) were also biased towards cell type-specific HOT loci and, conversely, that DFMs strongly associated with proximal HOT loci (ETS and SP family) were biased towards ubiquitously expressed genes (Table S12, Figure 4B). The distal, cell type-specific class of DFMs differed among cell types with the GATA, NFE2, and TBX1 family of DFMs prominent in K562 cells and IRF8 and SPI1 DFMs prominent in GM12878 cells (Figure S9C-D). These distal, cell type-specific DFMs have nearly all been implicated in the regulation and differentiation of their corresponding cell lineage (Alder et al. 2014; Ferreira et al. 2005; Di Tullio et al. 2017; Davies 2013; Wang et al. 2008; Iwasaki et al. 2005). HOT loci that were common to all three cell lines were enriched for close proximity to housekeeping genes involved in cellular metabolism of organic compounds (Table S13). Conversely, cell type-specific, HOT loci tended to neighbor corresponding cell type-specific genes (Figure S9E-G). In general, we found cell type-specific genes were more likely (49% vs. 16%, chi-square P<1E-6) to contain multiple, cell type-specific HOT loci within 50kb of their TSS, while ubiquitously expressed genes were more likely (59% vs. 9%, chi-square P<1E-6) to have an ubiquitously HOT promoter (Figure 4C). These data support a model of multiple, cell type-specific HOT loci bound by cell type-specific, driver DAPs regulating cell type-specific gene expression (Figure 4D) and ubiquitously expressed housekeeping genes regulated by an ubiquitously HOT promoter bound by common ETS or SP family DAPs (Figure 4E).

### Copy number variation, 3D chromatin structure, and GC content associate with HOT loci

To further explore mechanisms underlying the formation of HOT loci, we examined a variety of genomic characteristics linked to sites with high-densities of DAPs. In agreement with previous studies of TF motifs and flanking regions (Dror et al. 2015), we found HOT loci to be enriched for elevated GC content (rho=0.387, P<5E-16, Figure 5A). In addition to being a by-product of increased motif content, previous studies have proposed that elevated GC content, particularly in promoter regions, may lead to the formation of secondary DNA structures that induce indirect or non-specific DAP associations (Wreczycka et al. 2019). This hypothesis was difficult to test directly as both DAP recruitment and promoter GC content were associated with neighboring gene expression (rho=0.051, P<5E-16, Figure S10). However, we found gene expression levels to be an independent predictor of the number of promoter DAPs after correcting for promoter GC content (regression F-statistic P<5E-16) and found little variation in the strength of the correlation between gene expression level and number of promoter DAPs based on promoter GC content (Table S14, Figure 5B), which disfavors the idea that elevated GC content artificially drives the number of DAPs beyond gene expression-based expectations. There was no association between total repeat masked sequence, repetitive element composition, and locus mappability and the number of DAPs (Figure S10B-D).

**Figure 5.**
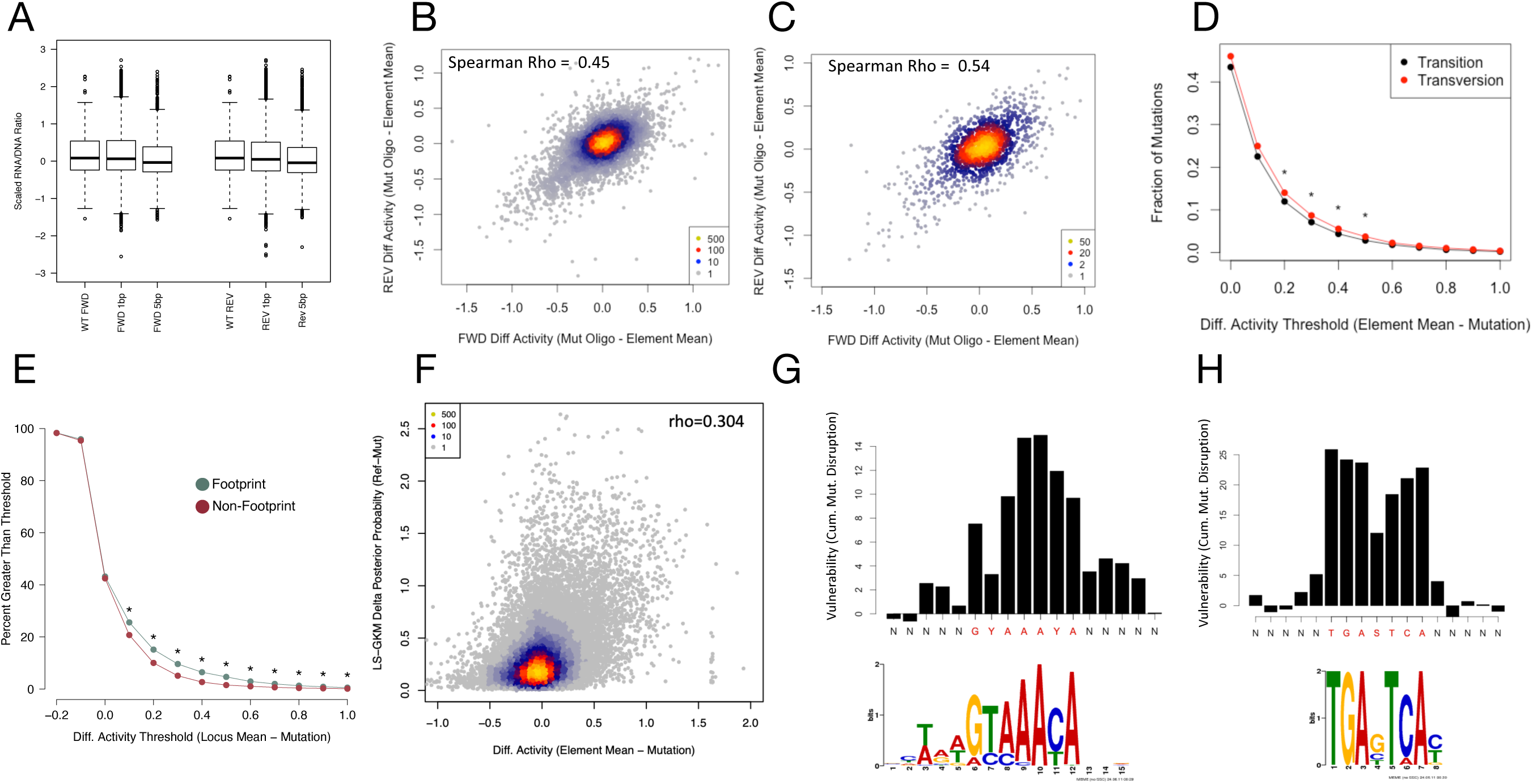
(A) Violin plots demonstrating the GC content of loci with increasing numbers of DAP ChIP-seq peaks. Width of each violin indicates the relative fraction of data contained. Boxes represent the median of each bin and whiskers are drawn to the 25^th^ and 75^th^ percentiles. (B) Scatter plot demonstrating the association between gene expression and DAP ChIP peaks in each genes promoter. Points and trend lines are colored based on promoter GC content. Promoters with GC content in the upper 50^th^ percentile of GC content (High) are colored green and those in the lower 50^th^ percentile of GC content (Low) are colored red. (C) Stacked bar plots showing proportion of loci with various levels of ChIP-derived DAP associations in genomic regions with heterozygous deletions, amplifications, or normal copy number. (D) Violin plots demonstrating the correlation between the number of ChIP-defined DAP associations and the number of Promoter Capture C interactions. Boxes represent the median of each bin and whiskers are drawn to the 90^th^ percentile. P-value reported is derived from Spearman rho correlation of the entire dataset. The sample size for each violin from left to right is 194,028, 37,084, and 13,792. (E) Boxplots demonstrating the fraction of DAPs in common between interacting loci and matched non-interacting loci for HepG2 Promoter Capture C. (F) Line plot indicating the relative fraction (cell type-specific/ubiquitously expressed) of gene promoters with at least the specified number of Promoter Capture C interactions with other HOT loci. Interactions with cell type-specific loci are shown in green and interactions with loci that are HOT in all three cell lines are shown in red. The gray shaded area represents the 95% confidence, null interval of randomly shuffled loci interactions between cell type-specific and ubiquitously expressed promoters. The 500 cell type-specific and expression-matched ubiquitously expressed genes were identical to those selected in Figure 4.

Another potential mechanism driving ChIP-seq signal inflation is chromosomal ploidy differences or smaller scale copy number variation (CNV). Increasing the number of available DAP binding sites by copy number amplification could provide greater opportunities for DAP recruitment, resulting in a proportionally greater number of DNA fragments as input to the ChIP-seq assay and improved sensitivity for DAPs that may be incompletely accounted for with genomic background controls (Zhang et al. 2008). A gross assessment of the chromosomal distribution of HOT loci in HepG2 suggests this is an important variable to consider (Figure S11A-C). Increased ploidy of chromosome 20 and partial chromosomal amplifications of chromosomes 16 and 17 have been previously described in HepG2, and we observed that chromosomes 16, 17, and 20 harbored more HOT loci than expected based on their size and gene density (López-Terrada et al. 2009). However, we did not observe a higher rate of HOT loci on chromosomes 2 and 14, which have also been described as having increased ploidy in HepG2, arguing that ploidy alone does not drive extreme numbers of DAP associations. Moreover, the K562 and GM12878 cell lines had much smaller chromosomal deviations in rates of HOT loci, despite having an equivalent number of total HOT loci (Figure S11B-C). Intersecting ENCODE copy number variation (CNV) array data with our merged ChIP-seq peak loci, we found a significant depletion in DAP associations at loci with a heterozygous deletion compared to loci with normal copy number (0.7% vs. 5.4%, Chi-squared P=6.9E-50, Figure 5C). There was only a minor enrichment for HOT loci in amplified regions relative to normal copy number regions (5.4% vs. 6.9%, Chi-squared P=2.9E-25) and 20% or less of loci at any DAP-association threshold were found in amplified regions (Figure S11D). Thus, locus copy number appears to be a statistically significant yet relatively minor contributor to the observed DAP association patterns.

Three-dimensional chromatin structure might also contribute to the observed pattern of DAP co-associations. The importance of 3D chromatin structure is becoming increasingly recognized and much of this structure is thought to be driven by large protein complex interactions with DNA (Quinodoz et al. 2018). Protein complexes that bring together multiple loci on the chromosome could give the appearance of dramatic levels of indirect ChIP-seq binding at each locus involved in a given network. Analysis of Promoter Capture C and Chromatin Interaction Analysis with Paired-End Tag (ChIA-PET) data recently generated in HepG2 cells revealed a weak positive correlation between the number of DAPs and the number of 3D interactions detected across loci (rho=0.236, P<5E-16, Figure 5D,S12A) (Chesi et al. 2019). Interacting loci did share a significantly higher proportion of DAPs than non-interacting loci (Figure 5E). In agreement with the model proposed in Figure 4D-E, cell type-specific promoters were significantly more likely to exhibit distal Capture C interactions with other cell type-specific HOT loci than promoters of ubiquitously expressed housekeeping genes. These association trends were also found by ChIA-PET and chromatin capture available for the K562 and GM12878 cell lines (Figure S12B-E), and restricting these analyses to ssTFs or nssDAPs did not dramatically alter the strength of these correlations (Table S8) (Mifsud et al. 2015). Overall, we found the association between 3D chromatin interactions and DAP density to be weak but consistent across cell lines, and it is possible that they operate more strongly at a minority of HOT loci. However, HOT loci tend to cluster near each other relative to loci with low numbers of DAPs (Figure S12F), leading us to expect that these associations will likely strengthen as experimental approaches for examining 3D chromatin mature.

## Discussion

We have performed an extensive analysis of DAPs across three cell lines using 647 ChIP-seq experiments and 941 *in vitro* derived motifs. In each cell line, we found ∼15,000 loci that harbored ChIP-seq peaks for more than 25% of DAPs assayed. The number of HOT loci defined by this criterion appears to be consistent regardless of the number of DAPs incorporated into our analysis. Thus, we believe this result will be generalizable to future analyses that will incorporate increasingly comprehensive databases of genome-wide DAP associations. However, until all expressed DAPs have been assayed in a given cell line, it will be difficult to appreciate the total number of DAPs capable of participating in a single locus. As the prevalence of ChIP-seq peaks was only loosely correlated with their corresponding DFMs at HOT loci, a substantial proportion of signal at HOT loci is likely to be driven by indirect binding not constrained by the presence of specific motifs. Thus, the true upper limit for the number of DAPs detectably associated with a given locus yet may exceed the number of DAPs currently assayed. Moreover, HOT loci identified in our analysis are distinct from previously blacklisted regions shown to be common high signal artifacts in sequencing assays and are present at a majority of active enhancers and promoters in the cell lines we analyzed. While it is extremely difficult to differentiate indirect DAP binding from non-specific or artifactual ChIP-seq signal previously proposed to contribute to HOT loci (Wreczycka et al. 2019), the pervasiveness of complex DAP co-associations in non-ChIP-seq dependent DFMs and the predictable nature of regulatory activity modulation by mutation of constituent motifs suggests these observations are likely not purely due to ChIP artifacts. Furthermore, regardless of underlying mechanism, we find the number of DAP co-associations to be a useful marker of active regulatory elements. Rather than being a rare event capable of being filtered from future experiments, these loci appear to be a defining mark of neighboring transcription.

We do not yet know the mechanism(s) driving the HOT DAP association pattern, in part because of technical limitations of the ChIP-seq assay. Most critically, robust ChIP-seq requires a population of cells as input. Thus, it is impossible to conclude from these data what proportion of DAPs simultaneously co-associate in the same cell. Single-cell ChIP-seq is still in its infancy, but as it matures, it may provide important clues to assist in answering this question (Rotem et al. 2015). Furthermore, the allele-specificity of DAPs was not considered by our analysis. Few allele-specific analyses have been conducted on a large number of DAPs in the same cell line or tissue, but some evidence exists that DAPs may favor a single allele in the context of allelic sequence variation (Ramaker et al. 2017; Reddy et al. 2012).

Our analysis found allele copy number to be a minor contributor to the number of observed DAPs at a locus. Specifically, heterozygous deletions were slightly depleted for high numbers of DAPs and amplified loci showed a small increase in HOT loci in terms of proportional representation. We hypothesize that this is largely a technical artifact due to imperfect baseline normalization at high copy number loci in the ChIP-seq assay. However, there may be selective pressures driving amplification of HOT loci in rapidly dividing cancer cell lines. Additionally, the 3D chromatin structure at a locus is correlated with observed DAP co-associations. In particular, HOT loci are enriched for greater numbers of 3D interactions and a greater number of shared DAPs are observed between equivalently bound interacting loci than non-interacting loci. These data coupled with the tendency of at least a subset of HOT loci to cluster near one another in the genome support a previously described long-range “flexible billboard” model of enhancer function (Vockley et al. 2017; Arnosti and Kulkarni 2005). This model proposes that enhancer output is largely dictated by the aggregate sum of interacting motif and “tethered,” non-motif driven DAPs, which have been shown to co-localize in high concentrations via phase-separated condensates (Shrinivas et al. 2019), rather than rigidly organized, enhanceosome structures.

High numbers of DAP associations, whether they represent direct or indirect DNA binding, do strongly predict regulatory activity. This finding is largely independent of any initial model or classification based on activity or local clustering (e.g. super-enhancers) (Whyte et al. 2013). We found that DAP densities were associated with independently defined promoter and enhancer annotations, motif constraint, neighboring gene expression level, and reporter assay activity. HOT loci, as defined by ChIP-seq, tended be evenly distributed across enhancers and promoters; however, loci particularly enriched for DFMs were more likely to be promoters than enhancers.

The number of DAPs at a locus exhibited a roughly continuous relationship with neighboring gene expression and the ability to drive expression with a minimal promoter in a reporter assay. We did not find a saturation point beyond which greater numbers of DAPs were redundant; thus we hypothesize that DAPs contribute to a locus’ regulatory activity in a dose-dependent manner. HOT loci tended to naturally dichotomize into cell type-specific or cell type-ubiquitous groups. Cell type-specific genes tended to possess multiple, distal neighboring cell type-specific HOT loci that typically contain cell type-specific “driver” motifs such as HNF4A or GATA. Conversely, universally expressed, housekeeping genes generally had a ubiquitously HOT promoter containing SP or ETS family motifs. Thus, loci that play a role in regulating cell maintenance and differentiation seem to be readily identifiable by high densities of DAPs and can be readily segregated based on their constituent motifs. Because DAPs tend to aggregate at HOT loci in a partially cell type-specific manner, it may be difficult to fully impute the locations of HOT loci in other cell lines that have not been as extensively assayed as the core ENCODE cell lines included in this study. However, we found DFM-defined HOT loci overlapped heavily with ChIP-defined HOT loci, which may obviate the need to perform extensive numbers of ChIP-seq experiments in every cell and tissue type to predict the presence of a HOT locus.

Lastly, our STARR-seq results provide high-resolution data on the most important sequence elements governing activity of hundreds of HOT loci. An important observation from our data is that a majority of regulatory activity can be localized to a central 130-bp region of maximal ChIP-seq peak signal at a given locus and that equivalently-sized flanking regions showed activity roughly equivalent to our null sequences. We also found that activity at HOT loci can be dramatically altered in a predictable manner by some single base pair mutations. We found that HOT loci were most vulnerable to SNVs in previously identified, highly conserved portions of their constitutive motifs. In particular, a subset of ssTF motifs, including HNF4A, SP1, SP5, ETV4, FOXA and JUN/AP-1 motifs were highly prevalent at HepG2 HOT loci and were particularly enriched for high-impact SNVs in our mutagenesis assay. We believe this provides sufficient evidence to nominate these ssTFs as putative drivers of regulatory activity at HOT loci, and future experiments specifically modulating the activity of these ssTFs or their motifs at HOT loci will be informative. We also found evidence that the total number of DFMs at a locus can reduce its overall vulnerability to SNVs, suggesting that at least some HOT loci may be buffered from the effects of otherwise harmful mutations. This phenomenon is also apparent in the reduced effect size of GTEx eQTL SNPs that map to HOT loci. We believe similar mutagenesis experiments would need to be performed on roughly an order of magnitude greater number of loci to definitively test this hypothesis; however, our results justify further exploration, as this buffering effect potentially complicates the interpretation of non-coding variation that is naïve to the presence of neighboring DAPs.

Future investigation and interpretation of ChIP-seq and related data-types, especially when done on a single DAP or a small number of DAPs, will hopefully benefit from the knowledge that extensive DAP co-associations at a significant number of functionally pertinent putative binding sites may be present. We intentionally structured our analysis within a framework that is generalizable and can act as a resource for nominating potentially interesting loci for future experiments.

## Methods

### ChIP-seq data processing

BED files containing ChIP-seq peak information for the K562 and GM12878 cell lines were obtained directly from the ENCODE data portal (www.encodeproject.org) via the file accession number listed in Table S1. BED files containing ChIP-seq peak information for the HepG2 cell line were generated by the Myers and Eric Mendenhall labs under a consistent protocol in accordance with ENCODE standards and can be obtained from the GEO database under the GSE104247 accession. To define ChIP-seq derived DAP co-associations, we collapsed all neighboring peaks into a minimal set of non-overlapping 2-kb loci and defined all peaks within a bin as “co-associated”. This minimal set of 2-kb loci containing all ChIP-seq peaks was generated independently for each cell line in two steps. First, all peaks from each of a cell line’s BED files were merged into a single BED file with the bedtools package (https://bedtools.readthedocs.io/en/latest/) *merge* function and with the maximum distance required for merging (or –d flag) set to 2000 bp. Resulting merged peak loci that were smaller than 2kb were redefined by adding or subtracting 1 kb from midpoint to expand them to 2kb. Merged peak loci that grew larger than 2 kb (5.4% of the total) were split into contiguous individual 2-kb bins using split points that intersected the fewest possible ChIP-seq peaks in the original, individual DAP BED files. The small number of individual DAP peaks that were split in this process were assigned to the bin to which >50% of the peak resided. The resultant set of 2kb loci can be found in Table S3, Table S5, and Table S6 for HepG2, GM12878, and K562, respectively. DAPs were assigned to classes based on previous definitions (Lambert et al. 2018). This bed file binning method can be reproduced using the “SMART_BED_MERGE” repository in in the github repository: https://github.com/rramaker/GenomeTools2020/.

### CIS-BP motif footprint processing

All DAP motif position weight matrices (PWMs) were downloaded from the CIS-BP database (http://cisbp.ccbr.utoronto.ca/bulk.php) on 04/02/2018. Only motifs derived from in vitro methods (SELEX, protein binding microarray, or B1H) were included in further analysis. Motifs assigned to DAPs that were unexpressed (0 reads aligned) in each cell line based on expression data available on the ENCODE portal (HepG2 accession numbers: ENCFF139ZPW, ENCFF255HPM, GM12878 accession numbers: ENCFF790RDA, ENCFF809AKQ, K562 accession numbers: ENCFF764ZIV, ENCFF489VUK) were excluded from further analysis. ENCODE DNase-seq raw FASTQs (paired-end 36 bp) of roughly equivalent size (HepG2 accession numbers: ENCFF002EQ-G,H,I,J,M,N,O,P) were downloaded from the ENCODE portal and processed using the Kundaje lab, ENCODE DNase-seq standard pipeline (https://github.com/kundajelab/atac_dnase_pipelines) with flags: -species hg19 -nth 32 -memory 250G -dnase_seq -auto_detect_adapter -nreads 15000000 -ENCODE3. Processed BAM files were merged and used as input for footprinting with PIQ under default settings (Sherwood et al. 2014). Only footprints called with a PIQ Purity (positive predictive value) greater than 0.9 were used for subsequent analysis. High confidence DHS footprints were binned into a minimal set of non-overlapping 2-kb loci as described for ChIP-seq peaks above. The resultant set of 2-kb loci can be found in Table S4. TomTom was used to identify related DAP motifs. Specifically, DAP motif pairs that possessed a significant (FDR<0.05) similarity score or that shared significant similarity to another motif were treated as one motif capable of recruiting multiple DAPs as specified (Gupta et al. 2007).

### Intersecting with annotations of interest

ChIP-seq peak and DHS footprint loci were intersected with a variety of other genome annotations using the bedtools *intersect* and *map* functions. In all cases, >50% of the locus was required to overlap with a given annotation to assign it to an annotation. A source bed file containing IDEAS regulatory annotations was obtained from http://main.genome-browser.bx.psu.edu/. Loci containing an “Enh” annotation were designated as “strong enhancers”, those containing an “EnhW” annotation as “weak enhancers”, those containing “Tss”, “TssW”, “TssF”, or “TssCtcf” as “promoters” in the source file. All other annotations were grouped into an “other” class (Zhang et al. 2016). Gene coordinates were obtained from the ensemble genome browser (http://useast.ensembl.org/index.html) gene transfer format grch37.75 file. Gene expression data was obtained in the form of raw count data from the ENCODE data portal (HepG2 accession numbers: ENCFF139ZPW, ENCFF255HPM, GM12878 accession numbers: ENCFF790RDA, ENCFF809AKQ, K562 accession numbers: ENCFF764ZIV, ENCFF489VUK). Reads were normalized to counts per million (CPM) and averaged across replicates. Cell type-specific genes were defined as those having a 4-fold greater FPKM in a given cell line of interest than either of the other two cell lines and having a FPKM value of at least two in the cell line of interest. Cell type-ubiquitous genes were defined as those with an FPKM greater than five in HepG2, K562, and GM12878. HepG2 reporter assay data was obtained from previously published work hosted at the GEO accession GSE83894 in the file GSE83894_ActivityRatios.tsv (Inoue et al. 2017). Replicate average activity from the “MT” and “WT” columns were used for our analysis. GM12878 High Resolution Dissection of Regulatory Assay (HiDRA) data was obtained from previously published work hosted at the GEO accession GSE104001 in the file GSE104001_HiDRA_counts_per_fragmentgroup.txt (Wang et al. 2018). Fragments with zero plasmid DNA reads in any replicate were removed prior to analysis. Subsequently, fragment reads were normalized by counts per million and replicate median log10(RNA/plasmid DNA) ratios were used for our analysis. Significant liver GTEx eQTL SNPs were downloaded with permission from GTEx download portal. Specifically, we obtained the “Liver_Analysis.snpgenes” file from the V6 data release that contains significant eQTL SNPs derived from liver tissue expression data. GERP scores were obtained from the UCSC genome browser under the “Comparative Genomics” group. Copy number variation data was obtained from the ENCODE data portal under the file accession ENCFF074XLG. Deletions and amplifications were assigned as designated in the fourth column. Promoter Capture C data for HepG2 was obtained from previously published work hosted in the Array Express database (https://www.ebi.ac.uk/arrayexpress/experiments/) under the accession E-MTAB-7144 (Chesi et al. 2019). A processed bed file of significant interactions at 4 DpnII fragment resolution was graciously provided upon request of the Grant lab. Bed regions greater than 10kb were removed and regions less than 10kb were expanded at their midpoint to 10kb prior to further analysis. POLR2A ChIA-PET BED files containing significant 3D interactions for K562 were obtained from the ENCODE data portal under the file accessions ENCFF001THW and ENCFF001TIC. POLR2A ChIA-PET BED files containing significant 3D interactions for HepG2 were made available upon request from the Yijun Ruan lab. For both ChIA-PET analyses, BED regions greater than 10kb were removed and regions less than 10kb were expanded at their midpoint to 10 kb prior to further analysis. Promoter capture Hi-C bed files for GM12878 was obtained from previously published work hosted in the Array Express database (https://www.ebi.ac.uk/arrayexpress/experiments/) under the accession E-MTAB-2323 (Mifsud et al. 2015). We used the TS5_GM12878_promoter-other_significant_interactions.txt file for analysis. Similar to POLR2A ChIA-PET data above, bed regions greater than 10kb were removed and regions less than 10kb were expanded at their midpoint to 10kb prior to further analysis. BED files containing repetitive element alignment scores were obtained from the UCSC table browser “repeat masker” track under the “Repeats” group. BED files containing DUKE 35mer mappability scores were obtained from the UCSC table browser “mappability” track under the “Mapping and Sequencing” group. All P values reported in the manuscript were capped at P<5E-16 to improve readability.

### STARR-seq library design and cloning

STARR-seq library consisted of 90,581 sequences representing 390 bp within 245 unique loci in both the forward and reverse orientation with tiled single base pair or 5-mer mutations. We selected loci that had previously demonstrated activity in the HepG2 cell line in Inoue et. al, 2017, because we reasoned a baseline level of reporter assay activity is required to see differential activity upon mutation (Inoue et al. 2017). Roughly two thirds of our loci met our specified HOT threshold all details regarding the loci assayed are available in Table S9. Alternate bases were randomly signed for single base pair mutations. 5-mer mutations were AAAAA or TTTTT depending on which was most divergent from the reference sequence. Previously demonstrated reporter activity in the top quartile of Inoue et. al. or in house data sets was the primary inclusion criteria (Inoue et al. 2017). Additionally, 50 negative control loci with low reporter assay activity based on previous in-house experiments and 371 GC-content matched control loci were included as negative controls. GC matched control sequences were generated using the *nullseq_generate* executable from the kmersvm website (http://beerlab.org/kmersvm/) on the provided hg19 genome indices (Fletez-Brant et al. 2013). Our complete oligonucleotide library is included in Table S15. Library oligonucleotides were synthesized by CustomArray as single stranded 170-bp sequences corresponding to 130 bp test elements (from either the 130 bp activity core, 130-bp left or 130-bp right flanking sequence for each locus) with 20-bp Illumina sequencing primer binding site tails. A first round of PCR was performed with the STARR-seq oligo amp F and R primers (Table S16) to amplify the library and generate double stranded DNA, complete the Illumina sequencing primer sequences and add 15 bp of sequence homologous to the hSTARR-seq (Addgene #99292) plasmid for InFusion cloning. PCR was performed with 20 reactions consisting of 1 uL CustomArray library input, 10 uM primers and the KAPA HiFi 2x PCR Master Mix (KAPA Biosytems) with the following conditions : 98 °C for 30 s, 20 cycles of 98°C for 15 s, 65°C for 30 s, 70°C for 30 s, and a final extension of 72 °C for 2 min. PCR products were pooled and cleaned up with the Zymo PCR Cleanup Kit (Zymo) before performing 2% agarose gel separation and extraction with the Zymo Gel Extraction Kit following manufacturer’s instructions. The cleaned up and amplified library was diluted to 10 ng/uL and 25 ng was used as insert in a 3:1 insert:vector InFusion reaction with 150 ng of hSTARR-seq plasmid (linearized with AgeI and SalI) in five replicate reactions following manufacturer’s instructions. InFusion reactions were pooled and cleaned and concentrated with 1.8X Ampure beads (Agencourt) in DNA LoBind tubes (Manufacturer) and eluted in 16 uL dH20. Six transformation reactions consisting of 2 uL of the cleaned and concentrated InFusion product were transformed into Lucigen Endura electrocompetent cells pooled and grown overnight at 37 °C in 2 L of LB ampicillin media at 200 RPM. Serial dilution plating of the transformation yielded an estimated library complexity of 9.7×10^7^ colonies, or roughly 1000x representation of the 9 x 10^4^ library elements. The full 2 L overnight culture was centrifuged at 5,500 RPM yielding a total pellet weight of 7.5 g from which the full plasmid library DNA was extracted using the Qiagen EndoFree GigaPrep and eluted in 2.5 mL TE buffer at a final concentration of 2.4 ug/uL following manufacturer’s instructions. Final library representation was determined by amplifying the insert library with the STARR-seq Sequencing Primers containing P5 and P7 Illumina sequences (Table S16) in 10 reactions consisting of 10 ng of plasmid library, 10 uM primers and the KAPA HiFi 2x PCR Master Mix (KAPA Biosystems) with the following conditions : 98°C for 45 s, 16 cycles of 98°C for 15 s, 65°C for 30 s, 70°C for 30 s, and a final extension of 72°C for 2 min. PCR products were pooled, run on 2 % agarose gel and extracted using the Zymo Gel Extraction Kit (Zymo) with final elution in 16 uL. The final sequencing library was quantified using Qubit dsDNA Broad Range (ThermoFisher) and the KAPA Library Quantification Kit (KAPA Biosytems) and sequenced on the Illumina MiSeq with PE 150 bp reads following standard protocols.

### STARR-seq library transfection, RNA isolation and library preparation

The STARR-seq library was transfected into HepG2 cells in 30 cm^2^ plates (25 million cells per plate) with 532 ug DNA using FuGene reagents at 4:1 ratio, with 12 replicate transfections. 24 hours after transfection, transfected cells were lysed on plate in RLT buffer (Qiagen) and stored at -80°C. The 12 replicates were condensed to 6 total replicates by combining cell lysates from two transfections. Total RNA was then isolated using the Norgen Total RNA Purification Kit using manufacturer’s instructions. STARR-seq libraries were prepared as previously described with any modifications included below (Gaulton et al. 2013). Poly-A RNA was isolated in triplicate for each replicate with 75 ug RNA input using Dynabead Oligo-dT_25_ beads (Life Technologies) with double selection and eluted in 40 uL 10 mM Tris-HCl. PolyA-RNA was then subjected to DNase digestion with TURBO DNase (Life Technologies) at 37°C for 30 min, cleaned up with the Zymo RNA Clean and Concentrate Kit (Zymo) and eluted in 50 uL RNA elution buffer. Reverse Transcription was performed with 45 uL of the cleaned and concentrated RNA for each replicate with 2 uM STARR-seq Gene Specific Primer (Table S16) and SuperScript III Reverse Transcriptase as previously described (Gaulton et al. 2013). cDNA from the RT was then treated with RNAse A for 1 hr at 37 °C, cleaned up with 1.8X Ampure beads and eluted in 100 uL Buffer EB. Junction PCR was performed in quintuplicate for each replicate with 20 uL cDNA and 10 uM input Forward and Reverse Junction Primers (Table S16) using KAPA HiFi 2x PCR Master Mix (KAPA Biosystems) with the following conditions: 98°C for 45 s, 15 cycles of 98°C for 15 s, 65°C for 30 s, 72°C for 70 s, and a final extension of 72°C for 1 min. Junction PCR cDNA reactions were pooled by replicate and cleaned up with Ampure beads and eluted in 100 uL dH20. After optimization, sequencing PCR was performed in quadruplicate for each replicate with 5 uL Junction PCR cDNA, 10 uM input STARR-Seq Sequencing Primer F and indexed Sequencing Primer R (Table S16) with KAPA HiFi 2x PCR Master Mix(KAPA Biosytems) with the following conditions : 98°C for 45 s, 5 cycles of 98°C for 15 s, 65°C for 30 s, 70°C for 30 s, and a final extension of 72°C for 1 min. Replicate sequencing PCR products were pooled, gel extracted with the Zymo Gel Extraction Kit (Zymo), eluted in 20 uL elution buffer and quantified with the Qubit Broad Range dsDNA kit. The six STARR-Seq RNA replicate libraries and a STARR-Seq Plasmid DNA input library (amplified as before but with 20 ng DNA input and 15 PCR cycles) were normalized with the KAPA Library Kit (KAPA Biosystems) and sequenced on an Illumina NextSeq with 150-bp paired-end reads using standard protocols.

### STARR-seq data processing and analysis

FASTQ files were adapter trimmed using cutadapt version 1.2.1 prior to alignment. Trimmed reads were mapped to our oligo library using bowtie 2 version 2.2.5. A custom bowtie index was generated with our oligo library (Table S15) in FASTA format using the build command under default settings. Trimmed FASTQ files were subsequently aligned to our custom index in a manner that required a perfect sequence match only in the correct orientation. Specifically, the --norc, --score-min ‘C,0,-1’, and -k 1 flags were used with the remainder under default settings. Aligned SAM files were converted to count tables using samtools version 1.2 indexing. This alignment procedure can be reproduced with the STARR_SEQ_Mutagenesis folder in the github repository: https://github.com/rramaker/GenomeTools2020. Our plasmid DNA library resulted in 49739461 reads with a 48.0% alignment rate. Our six RNA sequencing replicates resulted in an average of 31,942,788 reads (min = 4,189,978, max = 52,296,433). To balance read depths across replicates, we collapsed our six initial replicates into three replicates with an average of 63,885,576 reads (min = 56,486,411, max = 68,063,096) with an average alignment rate of 49.66%. A total of 90.4% of synthesized oligos were detected in our plasmid DNA library and 99.4% of test sequences were detected in at least one orientation. Ultimately, we applied a relatively strict count threshold filter by excluding oligos with less than 2 counts per million reads in our plasmid DNA library (44.7% of total oligos, 15.9% in both orientations) from further analysis.

Oligo activity was defined as replicate median log10(RNA CPM/DNA CPM). The differential activity of a mutation containing oligo, or the effect of a mutation on a locus, was computed as the difference in the mean activity of all oligos associated with a locus from the activity of a given mutated oligo of interest. In all cases the oligos containing forward strand sequence were analyzed separately from oligos containing reverse strand sequence for each locus. Raw count data and processed activity levels are available in Tables S17-18.

Predicted mutation effects were determined using the lsgkm analysis suite in a manner previously described (Lee 2016; Ramaker et al. 2017). Briefly, genome sequence was obtained in FASTA format for each HepG2 DAP narrow peak using the bedtools *getfasta* command. A GC content matched set of null peak sites 10 times greater in number than the number of peak observed for each factor was generated using the *nullseq_generate* executable from the kmersvm website on the provided hg19 genome indices as described above. SVMs were trained on narrow peak and matched background sequences using gapped 10-bp kmers and allowing for 3 non-informative bases using the “gkmtrain” executable obtained from the ls-gkm github webpage (https://github.com/Dongwon-Lee/lsgkm). All other settings were left at default. This resulted in 208 SVMs, one for each DAP analyzed in HepG2. Each mutant oligo sequence was scored with the SVM trained on each DAP and its resultant classifier value was subtracted from the reference sequence classifier value to determine a mutation’s predicted effect.

DAP enrichment for high impact mutations was computed using high confidence DHS motifs identified as described above. Enrichment P-values were calculated using a Fisher’s exact test comparing the ratio of high effect mutations (differential activity>0.25) to the total number of mutations falling within vs. outside of a given DAPs footprints.

### STARR-seq high impact mutation validation

To validate 14 identified SNVs with high impact in our assay, for each SNV we ordered ssDNA ultramers (Integrated DNA Technologies) corresponding to that SNV’s reference sequence, the high impact SNV sequence, and a neighboring low impact SNV, flanked by 15 bp primer binding tails (Table S19) for each SNV. To generate dsDNA suitable for inFusion cloning, we amplified Ultramers with primers containing sequence homologous to either the STARR-seq luciferase validation vector_mP_empty (Addgene# 99298) or pGL4.23 (Promega) (Table S16). Sequences were cloned into both vectors using inFusion cloning according to manufacturer’s instructions. Plasmid DNA was extracted from three separate colonies with the Spin Miniprep Kit (Qiagen) and sequence verified with Sanger sequencing (MCLAB, San Francisco, CA). Each colony was treated as a separate biological replicate for a given sequence as previously described (Whitfield et al. 2012). HepG2 cells were seeded at 40,000 cells per well in antibiotic free DMEM with 10% FBS in a 96-well plate. After 24 hours, 300ng of plasmid DNA for each biological replicate was transfected into HepG2 cells using FuGENE (Promega) in triplicate, resulting in 9 total replicates (3 biological X 3 technical) per sequence. Luciferase activity was measured 48-hours post-transfection with a 2-second integration time on a LMax II 384 Luminometer (Molecular Devices). Background subtracted luminescence values for each SNV were z-scored. Significance in expression was determined using a 2-tailed Student’s T-test.

### Data Availability

Raw FASTQs and processed count tables for our mutagenesis assay are freely available at the GEO accession number GSE142566. Please contact the first or corresponding authors for other intermediate files. Scripts required to reproduce the most computationally intensive portions of our analysis along with example test files are available at https://github.com/rramaker/GenomeTools2020.

## Acknowledgments

We are especially grateful to the Yijun Ruan and Struan Grant labs for uniform processing of the ENCODE ChIA-PET and Promoter Capture C data respectively. We also thank Eric Mendenhall and Surya Chhetri for their assistance with the alignment and quality control analysis of ChIP-seq experiments in HepG2, and are particularly grateful to them and the Myers/Mendenhall ENCODE group members, including Mark Mackiewicz, Kim Newberry, Dianna Moore, Laurel Brandsmeier, Sarah Meadows, Megan McEown, for generating the high-quality ChIP-seq data used in this paper. This work was supported by National Institute of Health (NIH) grants U54 HG006998-0 (to R.M.M. and E. Mendenhall) and 5T32GM008361-21 (to R.C.R. and A.A.H.).

## Supplemental Figures

**Figure S1.**
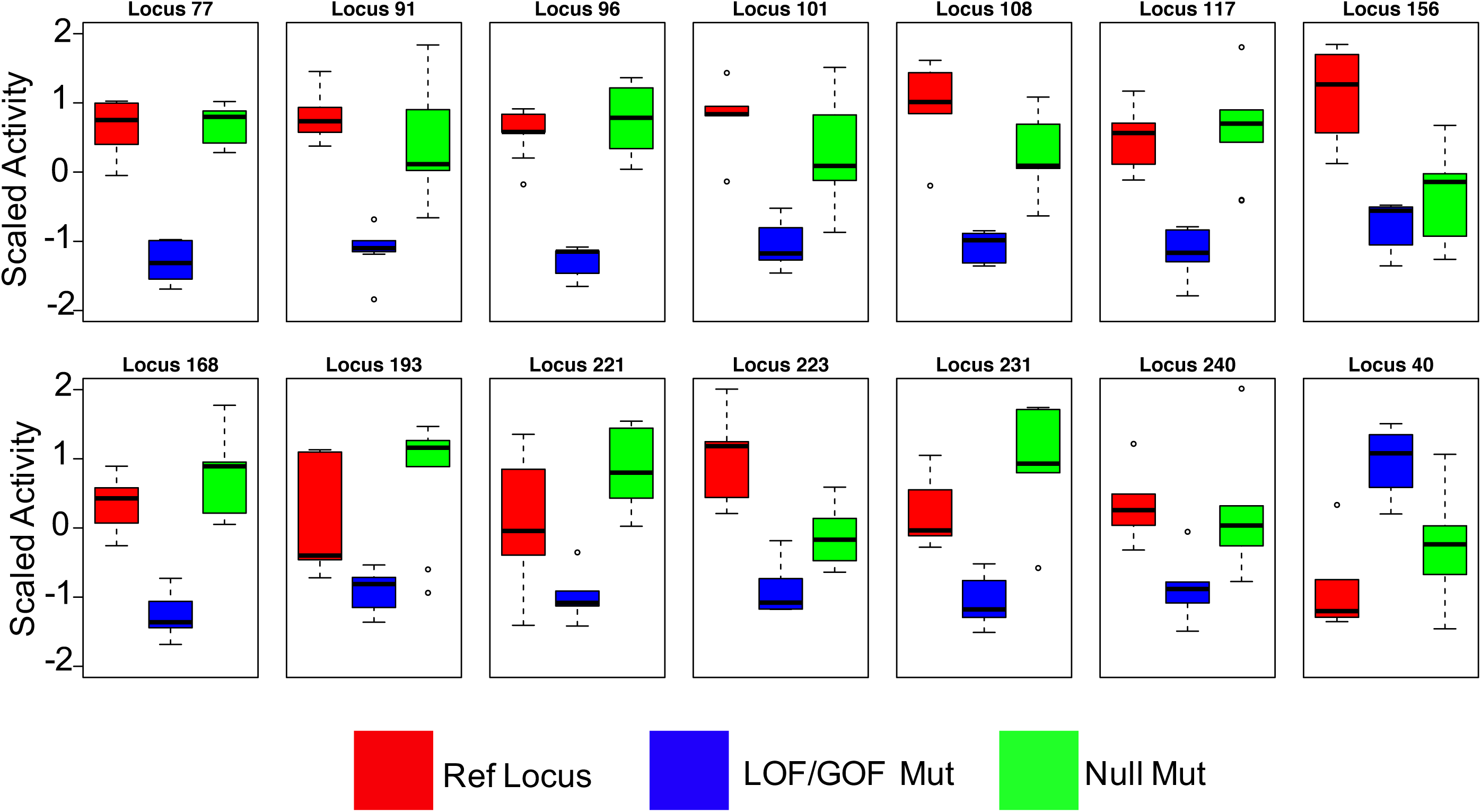
(A) Cumulative distribution function (CDF) showing the proportion of loci containing at least a given level of unique DAP or ssTF ChIP-seq peaks across the HepG2, K562 and GM12878 cell lines. Colors correspond to cell lines. Dashed lines indicate ssTF data only and solid lines represent data that includes all DAPs. (B) Number of loci reaching “HOT” threshold via the ChIP-seq assay after performing random down sampling of the number of DAPs included. Each data point for both plots represents the median result of a 100 random samples at a number indicated by the x-axis. The color of each line indicates the % of DAPs required to reach the ‘HOT’ threshold. (C) The recall performance or fraction of true HOT sites, as defined by the full dataset, detected with current sample of DAPs. Each data point for both plots represents the median result of a 100 random samples at a number indicated by the x-axis. The color of each line indicates the % of DAPs required to reach the ‘HOT’ threshold. (D-F) Number of loci reaching “HOT” threshold of 25% of DAPs associated via the ChIP-seq assay after performing random down sampling of the number of DAPs included. Each data point represents the result of a random sampling of a specified number of DAPs. The color indicates recall performance or the percentage of true HOT sites, as defined by >25% of DAPs bound in the full dataset, detected with current sample of DAPs. The black line represents the median result of 100 random samples of each number of DAPs as specified by the x-axis. Data for HepG2 (D), K562 (E), and GM12878 (F) is shown. (G) Cumulative distribution function (CDF) showing the proportion of loci included over an increasing exclusion threshold. The solid line represents observed HepG2 DAP ChIP-seq distributions and the dashed line represents a random shuffling of DAP assignments across all loci containing a peak. Random shuffling was performed in a manner that preserved the total number of DAP associations across all loci. (H) Venn diagram showing the overlap between HOT loci and super enhancers defined via the ROSE method. (I) Pie chart showing the number of DAPs associated with super enhancers. (J) Bar plot displaying the fraction of all HOT (black) or <10 DAP (gray) associated loci that have a specified combination of DFMs present.

**Figure S2.**
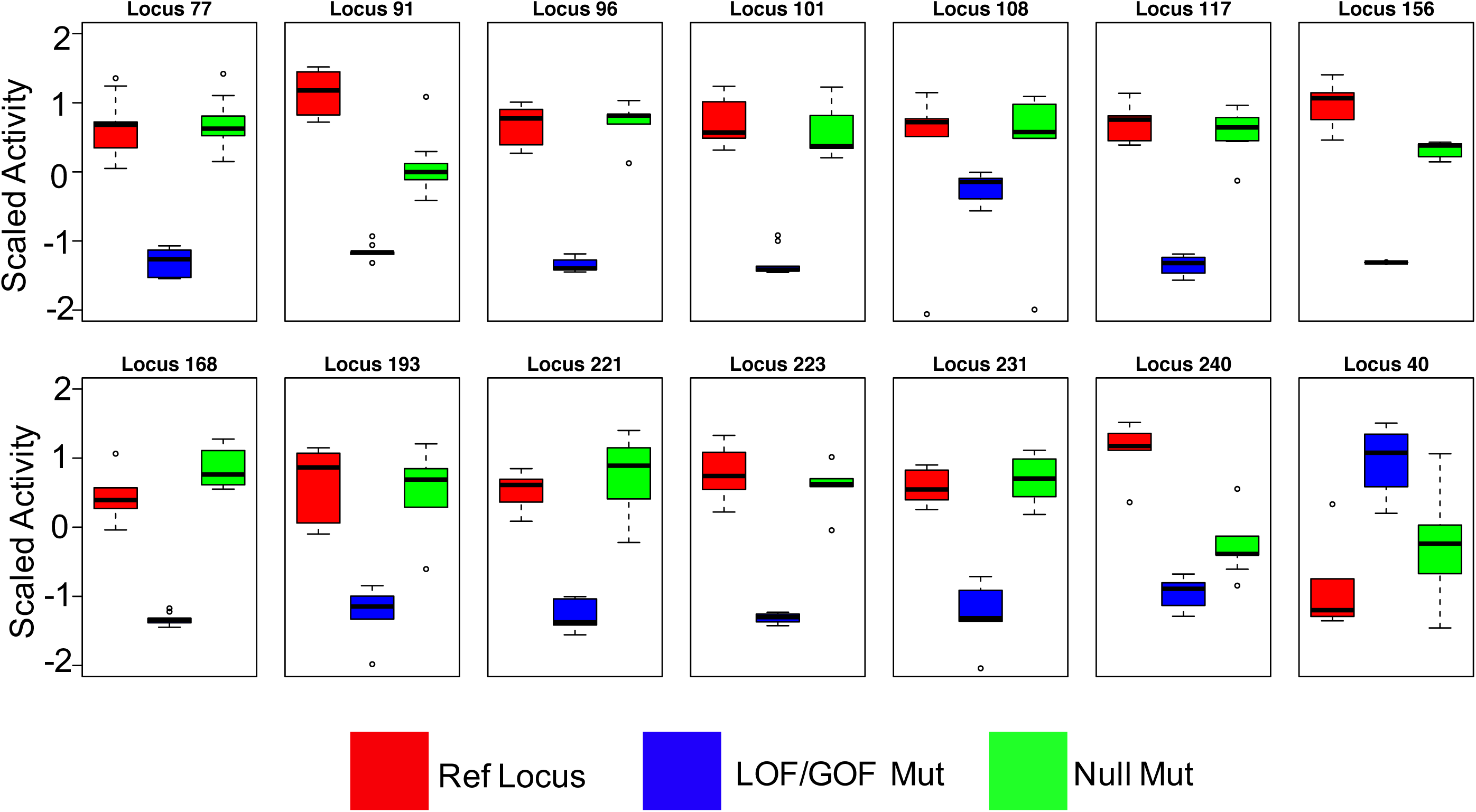
(A) IDEAS annotations of loci binned by the number DFM-defined DAP associations. Promoter, strong enhancer, and weak enhancer annotations represent 0.27%, 0.35%, and 0.22% of the HepG2 genome, while the remaining 99.16% of the genome (largely consisting of quiescent and repressed annotations) was used for the “Other” annotation. (B) Pie charts demonstrating the proportion of loci associated, based on ChIP-seq, with a specified number of DAPs for a variety of IDEAS annotations. (C-D) Histograms displaying the distribution correlations observed from 1000 random shuffles of gene-locus associations relative to that observed in the non-shuffled data sets shown in Figure 2B-C. (E-F) The expression level of the nearest gene to each loci binned by DFM-define DAP occupancy. Plots show loci distal (>5 kb, E) or proximal (<5 kb, F) to their nearest gene. (G) Stacked bar plots displaying the proportion of genes meeting a specified expression level bin with a specified number of neighboring HOT loci at a specified distance to TSS threshold. (H) GM12878 ChIP-defined DAP associations correlate strongly with previously published ATAC-STARR-seq reporter assay activity. Green boxes for plots D and E show loci binned by the presence or absence of common markers of enhancer or promoter regions that delineated a subset of the data.

**Figure S3.**
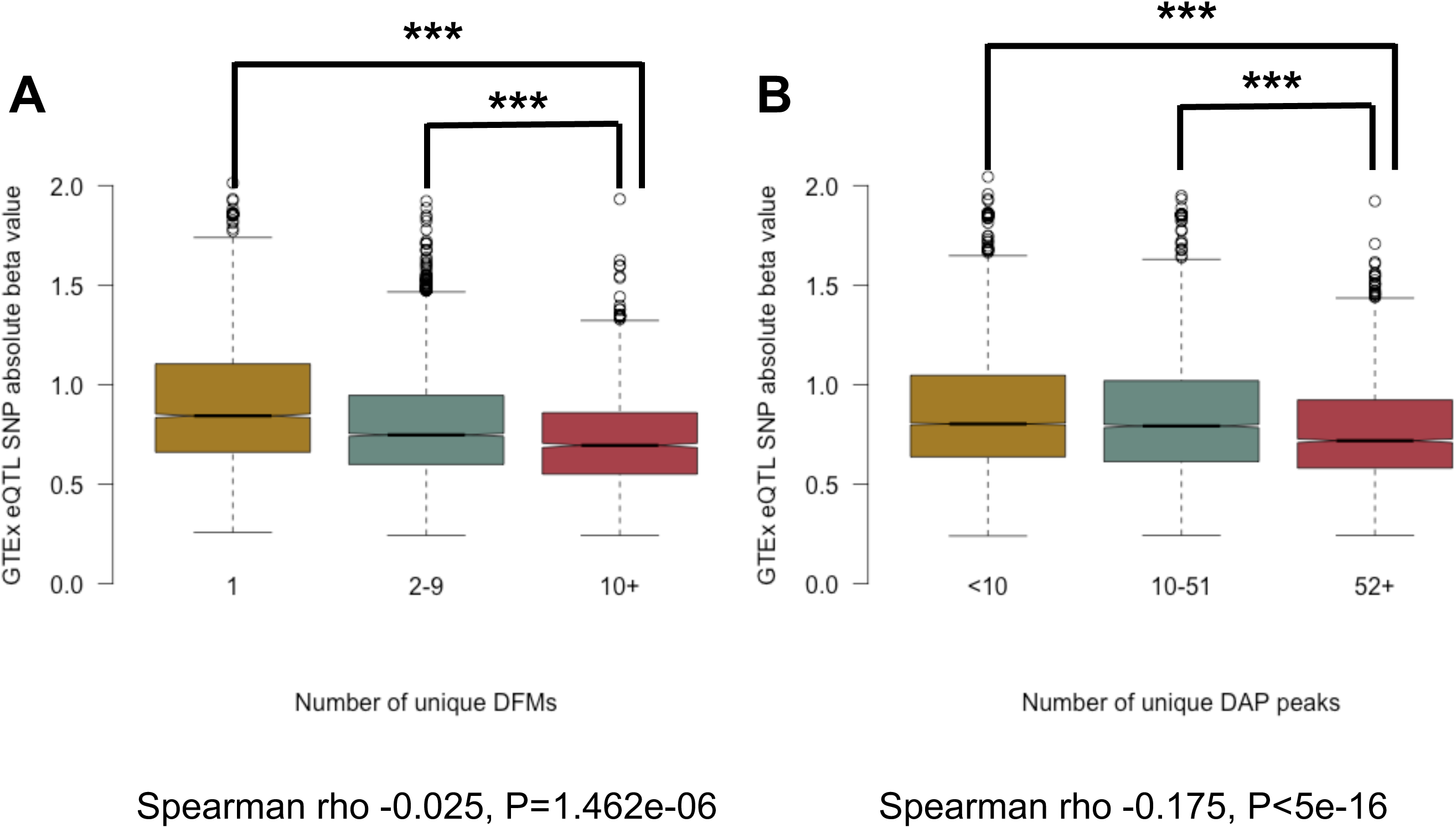
(A) Base-wise conservation at each position of the HNF4A motif as defined by GERP is plotted for each occurrence of the HNF4A motif with greater or less than 9 neighboring DFMs. (B-C) Spearman rho values representing the correlation between the median GERP score of each ssTF’s DFM and the number of unique DAP ChIP-seq peaks (B) or neighboring DFMs (C) in its 2kB locus. ***, **, and * denote Wilcox P-values of <0.05, <0.005, and <0.0005 respectively.

**Figure S4.**
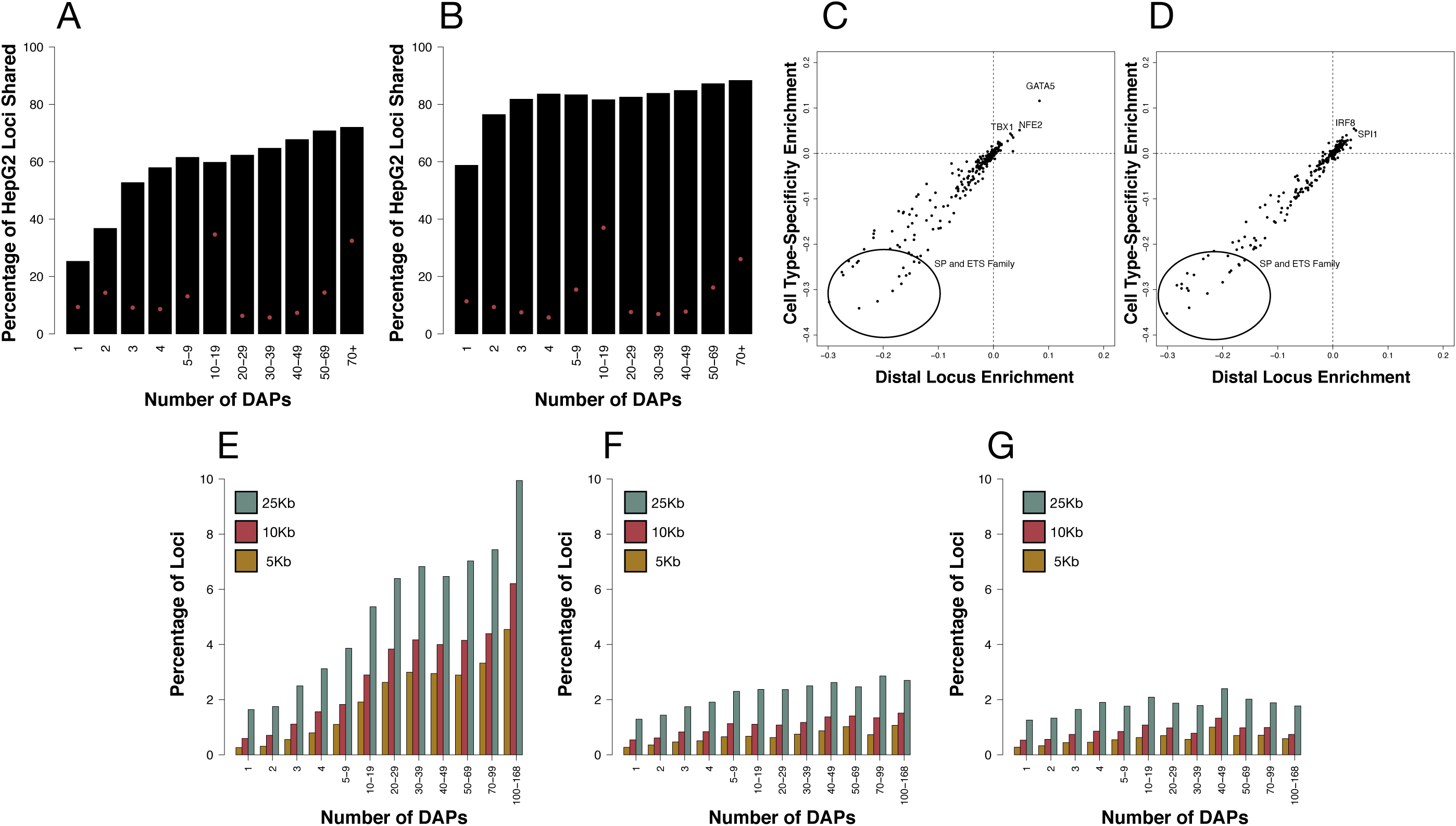
(A) Line plot indicating the positions of ChIP-seq peak (black) and DFM (red) pile up relative to point of maximum ChIP-seq peak pile up. Each point on the line indicates the average overlap, as a percentage of the maximum point of overlap, for all 13,792 HOT loci in HepG2. (B-C) Histograms indicating the distribution of reads assigned to oligonucleotides detected in DNA harvested from transfected cells. (B) represents data for all oligonucleotides including mutations. (C) represents only reference oligos from the “core” central 130-bp window of each assayed loci. (D) Boxplots demonstrating the number of counts for each single bp mutation oligo in the DNA input library. Each box contains 490 data points (one for each of our 245 loci in both orientations). A similar pattern was observed for our 5mer mutations suggesting no significant bias was introduced in our alignment methodology. (E-F) Scatter plot of the correlation between RNA counts per million (median across the 3 reps) and DNA counts per million. Plots are data subsets: (E) shows all reference HOT and null sequences, (F) shows central region reference elements and corresponding 1bp mutations, (G) shows central region reference elements and corresponding 5bp mutations, and (H) shows flanking reference elements and corresponding 5bp mutations. (I) Boxplots showing the fraction of oligonucleotides retained in our library at increasing DNA input representation thresholds. (J) Box plots showing replicate correlation of the RNA/DNA ratio at increasing DNA input representation thresholds.

**Figure S5.**
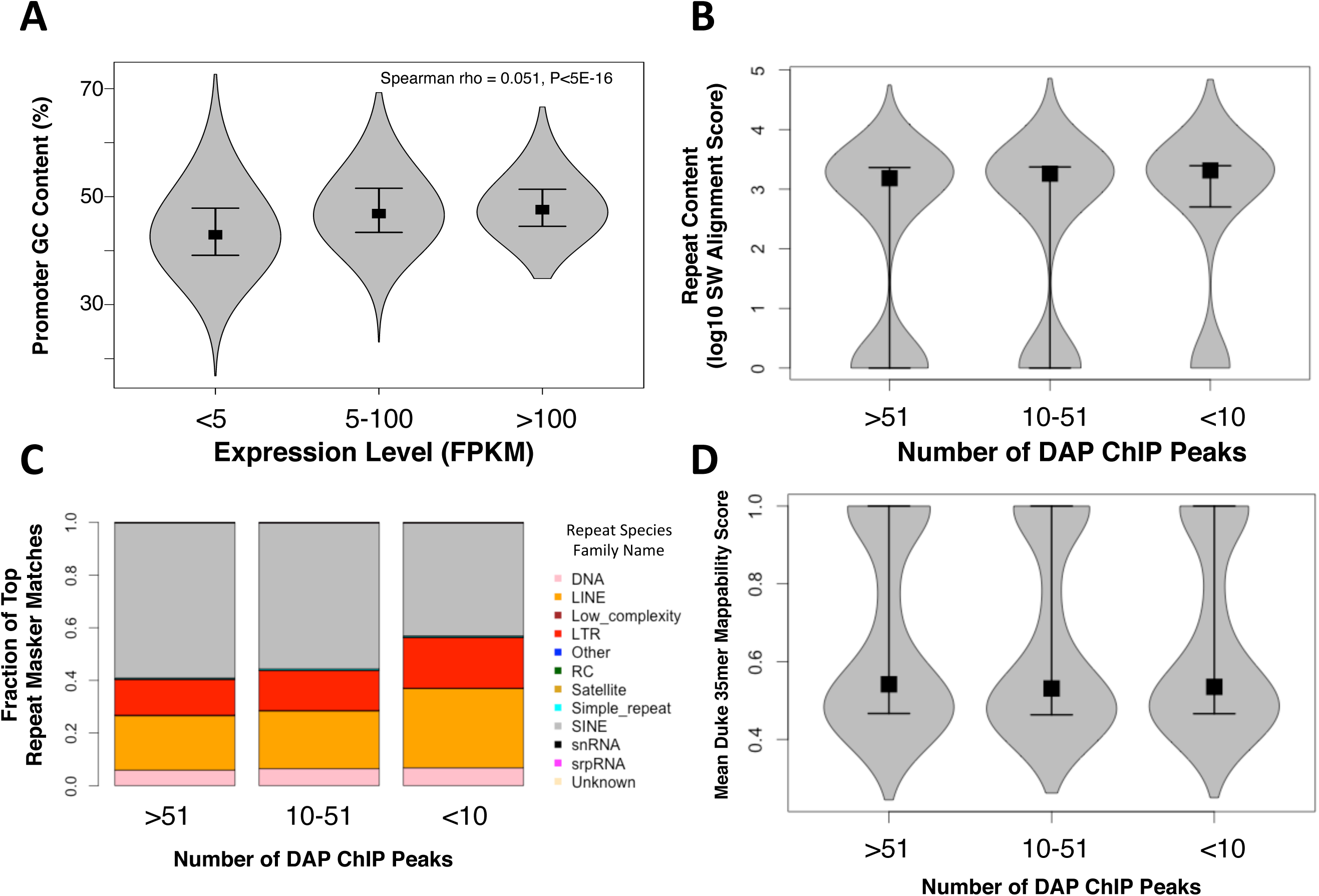
(A) Boxplots indicating that the distribution of activity in single mutant loci was roughly equivalent to that of WT regions with a slight decrease (Wilcox P<1E-16) in activity observed in 5-bp mutations. The left group of boxes shows data from the forward strand and the right group shows data from reverse strand oligonucleotides. (B-C) Scatter plots showing the correlation in the differential activity (mutant oligo – the locus mean) between the forward and reverse strands for single base pair mutations (B) and 5-bp mutations (C). The color indicates the number of points in contact with a given location on the plot. (D) Line plots showing the fraction of mutations that meet increasing thresholds of differential activity (loci mean – mutant oligo) for transitions versus transversions. (*) Indicates Fisher’s P<0.05. (E) Line plot indicating the proportion of mutations imposing a change of activity at a variety of thresholds for single base pair mutations. Green points indicate data for mutations falling within DHS footprints. Red points indicate data for mutations falling outside of DHS footprints. (*) Indicates Fisher’s P<0.05. (F) Scatter plot describing the correlation between LSGKM-predicted, DAP disruptions and differential activity of mutant oligos. Color indicates the number of points in contact with a give region of the plot. (G-H) Bar plots showing the cumulative differential activity (loci mean – mutation) across all positions in the FOXA (G) or AP1 (H) motifs. Activity correlates with the motif consistency at each position.

**Figure S6.**
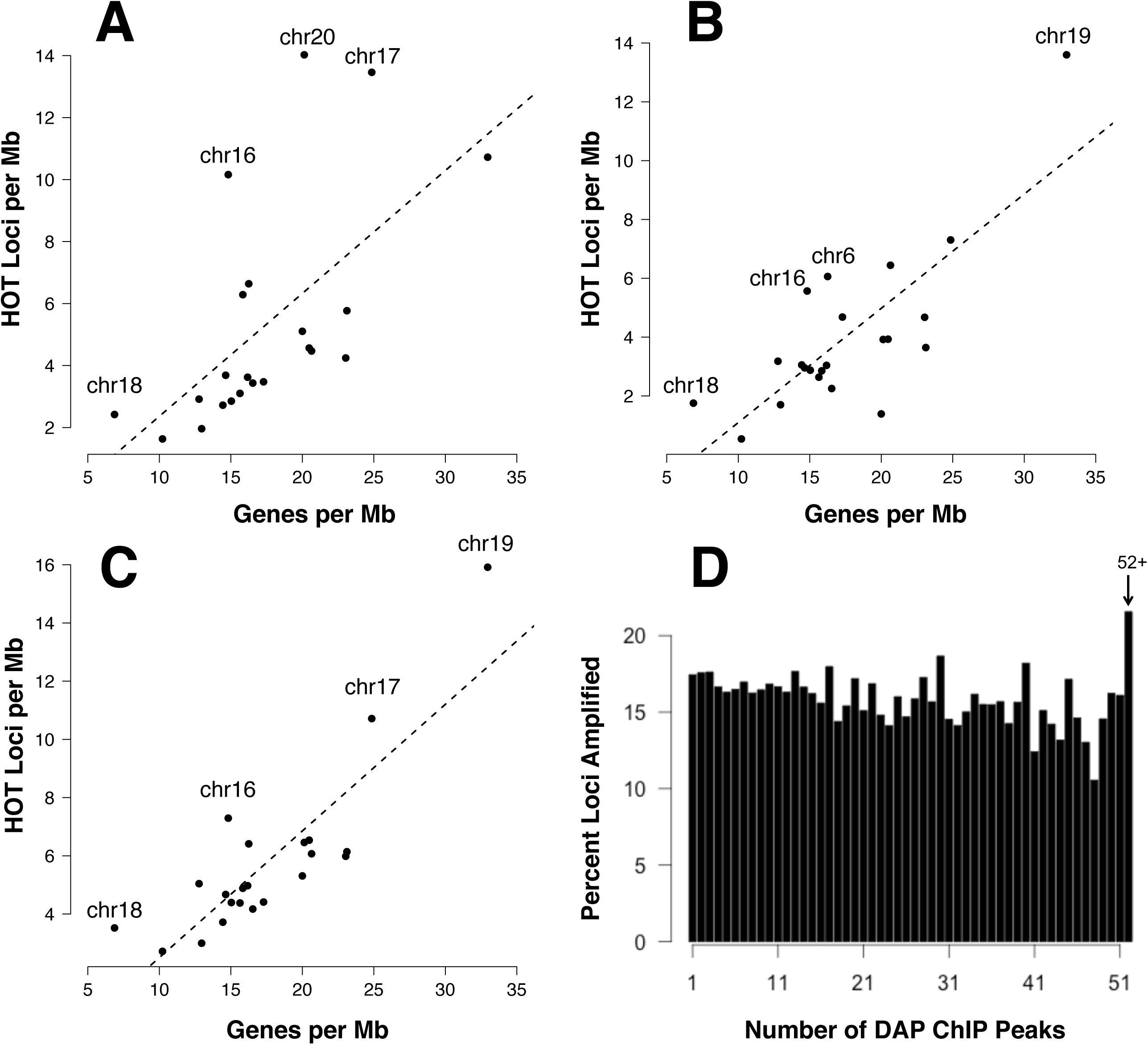
Validation data for the luc_mp_empty vector. Boxplots show inter-element z-scored data for reference (WT) oligos, oligos with loss of function (LOF) or gain of function (GOF) mutations, and oligos with predicted null mutations. Locus 40 is the only locus with a predicted GOF mutation. The remainder are predicted LOF.

**Figure S7.**
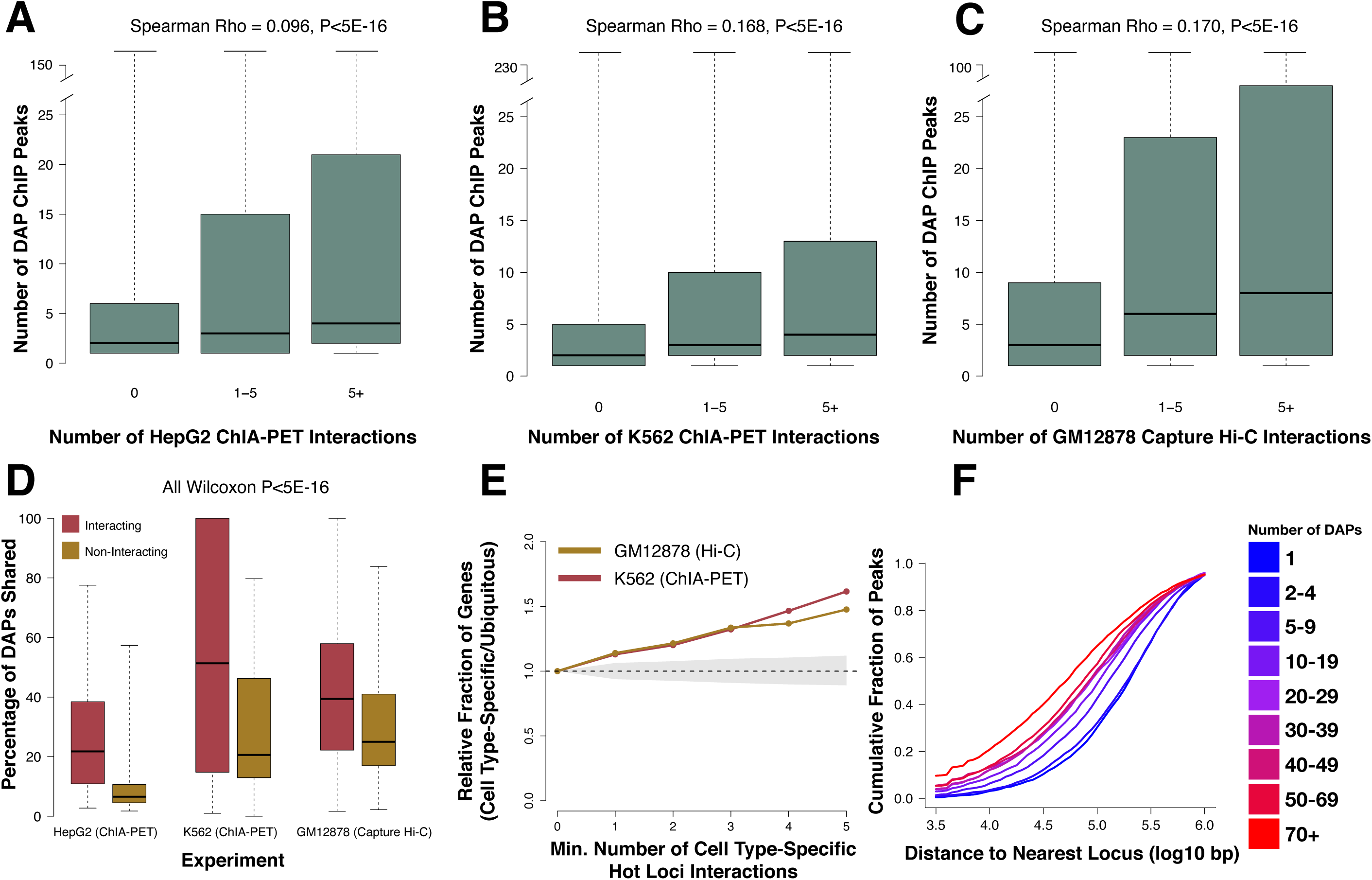
Validation data for the PGL4.23 vector. Boxplots showing inter-element z-scored data for reference oligos, oligos with loss of function (LOF) or gain of function (GOF) mutations, and oligos with predicted null mutations. Locus 40 is the only locus with a predicted GOF mutation. The remainder are predicted LOF.

**Figure S8.**
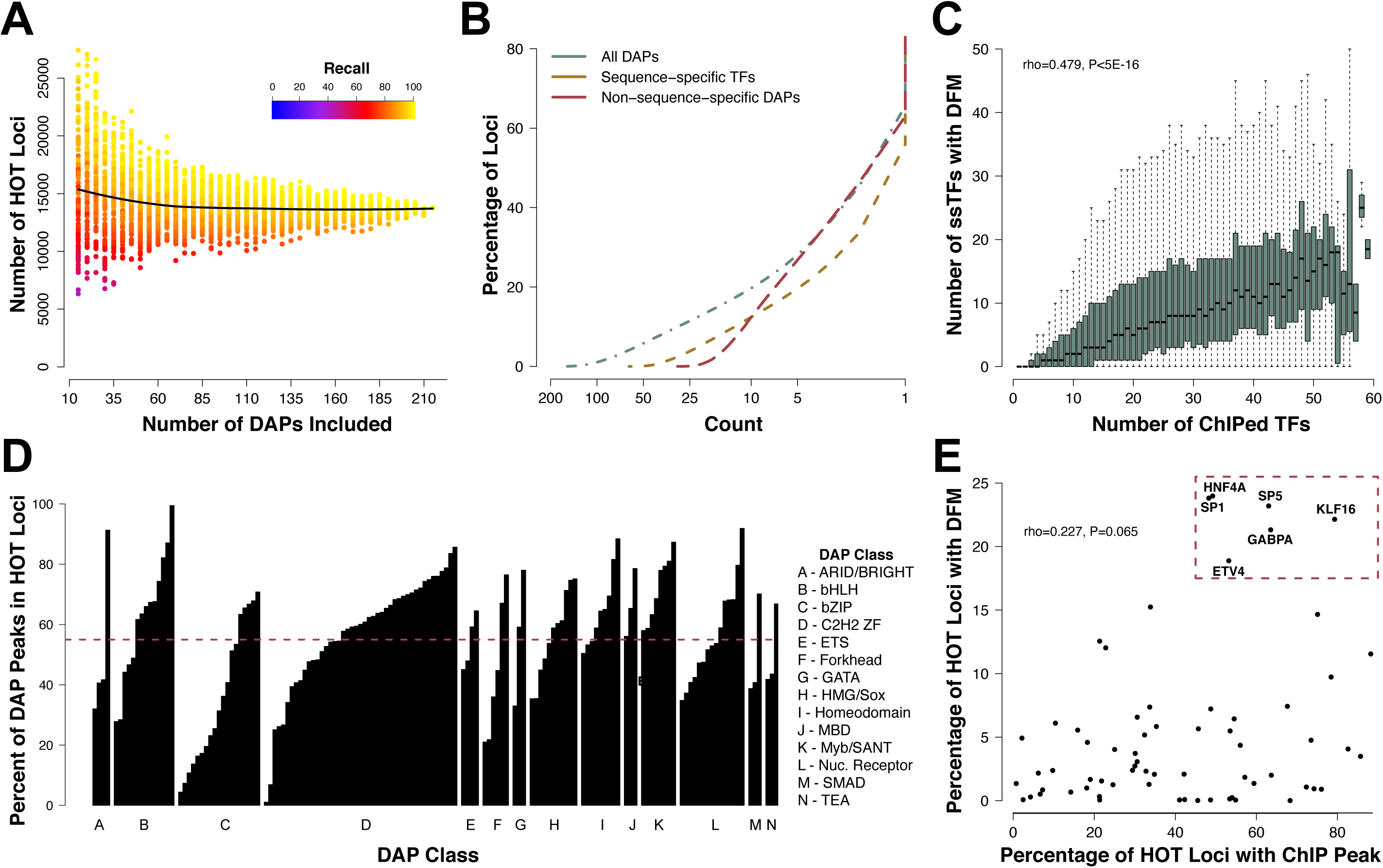
Boxplots indicating the magnitude of the effect size (absolute beta value) for significant (FDR<0.05) liver GTEx eQTL SNPs as function of the number of unique DFMs (A) or DAP ChIP-seq peaks (B) detected at the locus to which they map. *** Denotes Wilcoxon P<5e-16

**Figure S9.**
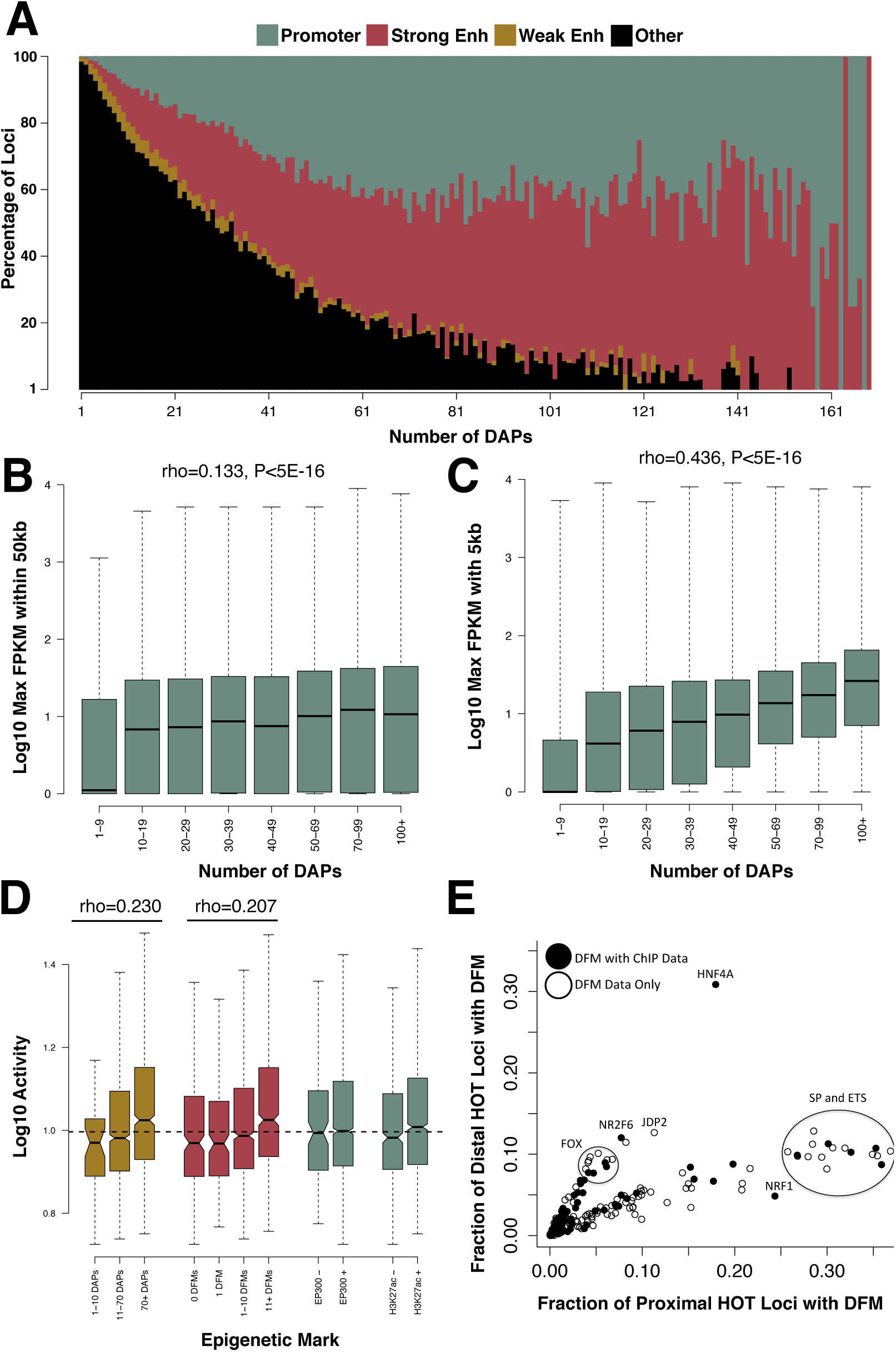
(A-B) The number of HepG2 loci bound by at least one DAP in GM12878 (A) and K562 (B). The red dot indicates the number of loci shared by an equivalent number of DAPs in both cell lines. (C-D) Scatter plot demonstrating the association between cell type-specific, HOT loci enrichment and distal, HOT loci enrichment in K562 (C) and GM12878 (D). Cell type-specificity enrichment value is computed by subtracting the fraction of cell type-specific HOT loci in which a DFM is present from the fraction of non-cell type-specific HOT loci in which a DFM is present. The distal locus enrichment is computed by subtracting the fraction of HOT loci >5kb from the nearest TSS in which a DFM is present from the fraction of HOT loci <10kb from the nearest TSS in which a DFM is present (E-G) Barplots indicating the fraction of each loci, stratified by number of ChIP-defined DAP associations in HepG2, that are present near HepG2 (E), GM12878 (G), or K562 (H) specific genes. Cell specific genes were computed by randomly sampling 500 genes that were expressed at least 4 fold higher in the cell line of interest than the other two cell lines and had an FPKM of at least five in the cell line of interest.

**Figure S10.**
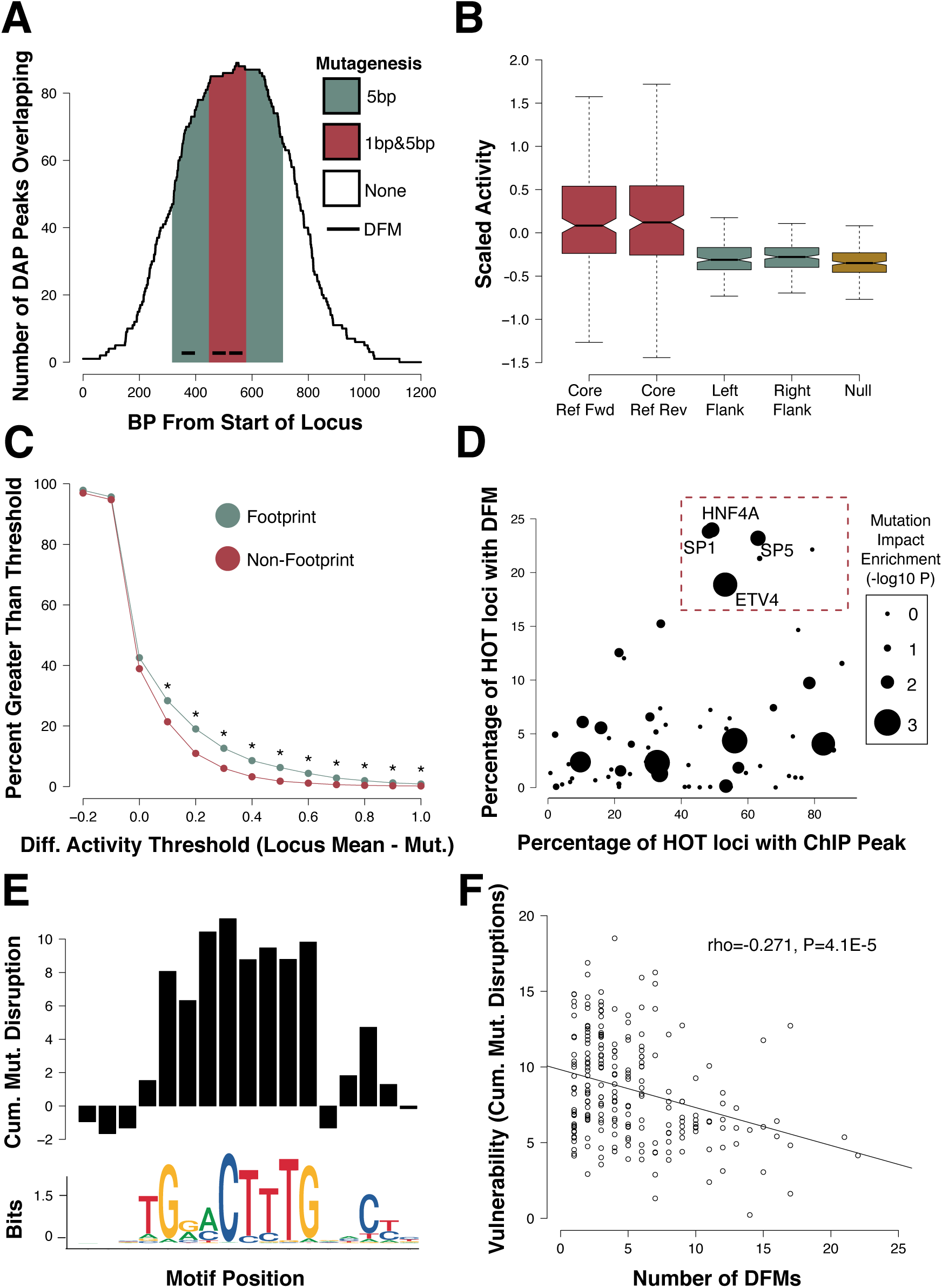
(A) Violin plot indicating percent GC content at promoters of genes with specified expression levels. (B) Violin plot displaying the distribution of top repeat masker Smith-Waterman alignment scores for loci with differing numbers of DAP associations. (C) Stacked barplots showing the family of repeat elements commonly found across loci. (D) Violin plots showing the distribution of Duke 35mer mappability uniqueness scores for loci with differing numbers of DAP associations. Violin plot contains a square at the median value and whiskers at 25^th^ and 75^th^ percentiles. Each violin contains data for 10000 randomly sampled loci meeting the “number of DAPs associated” criteria.

**Figure S11.**
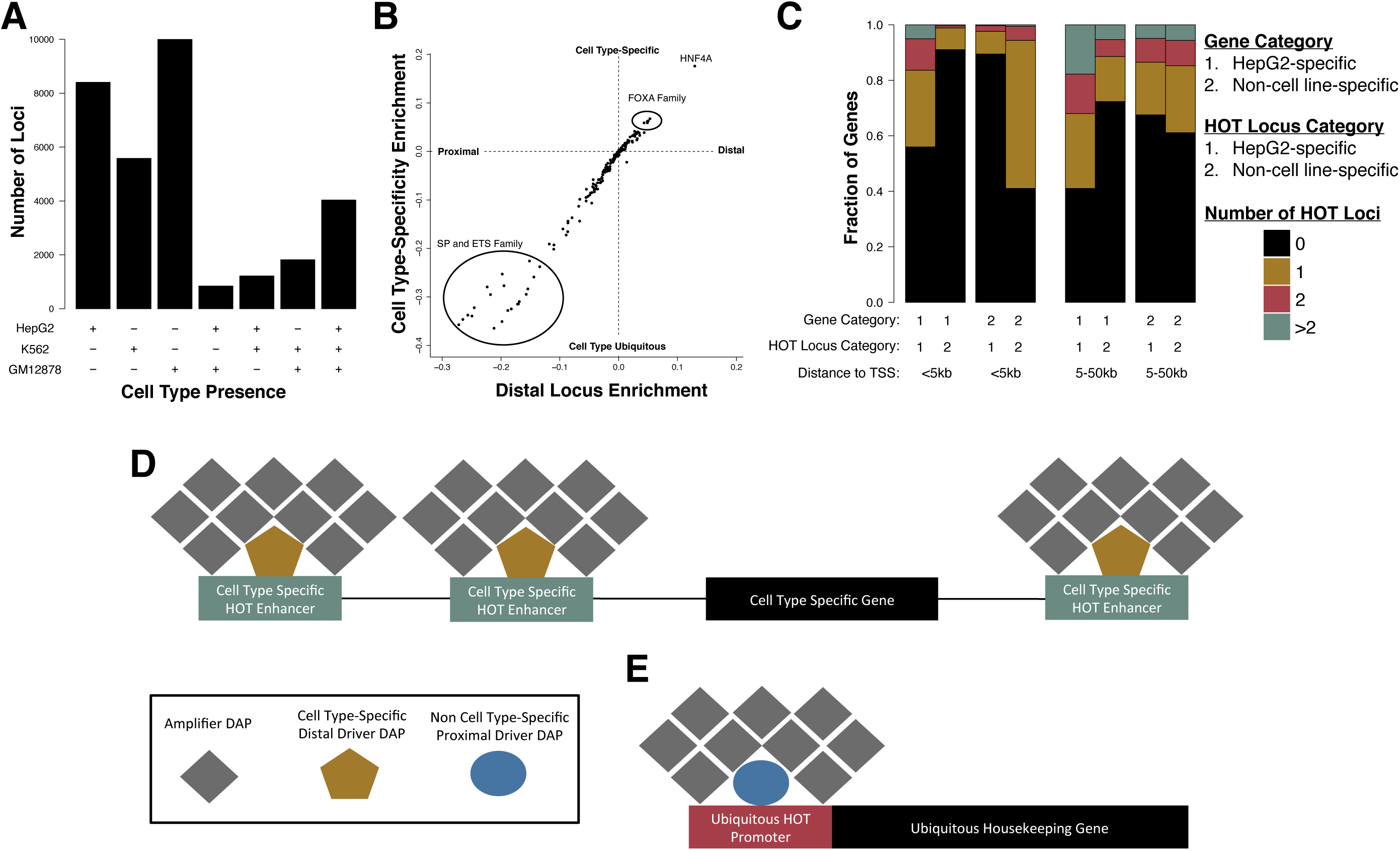
(A-C) Scatter plot indicating the observed rate of HOT loci vs. the expected rate based on gene density in each HepG2 (A), K562 (B), or GM12878 (C) chromosome. (D) Bar plots showing the proportion of loci amplified in HepG2 at increasing numbers of ChIP-seq derived DAP associations.

**Figure S12.**
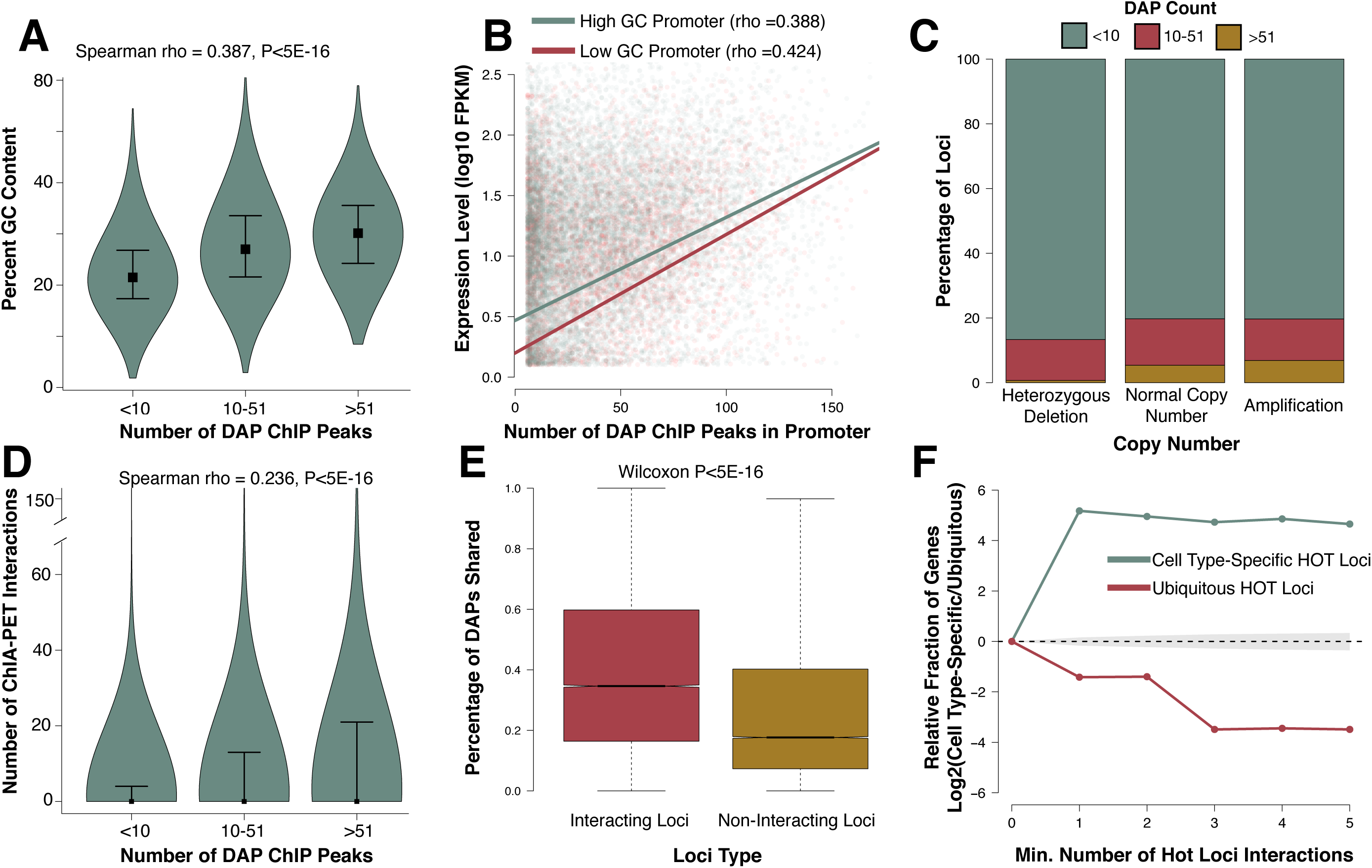
(A-B) Boxplots demonstrating the correlation between the number of DAPs bound (based on ChIP-seq peaks) and the number of ENCODE POLR2A ChIA-PET interactions observed in HepG2 (A) or K562 (B). The middle and high bins are divided at 5 interactions so roughly half of the non-zero loci are in each bin, in both cases. (C) Boxplots describing the correlation between the number of DAPs bound (based on ChIP-seq peaks) and the number of observed promoter capture Hi-C interactions in GM12878. P-values reported are derived from Spearman rho correlation of the entire dataset. (D) Boxplots demonstrating the fraction of DAPs in common between interacting loci matched non-interacting loci for HepG2 and K562 ChIA-PET data and GM12878 promoter capture Hi-C data. (E) Line plot indicating the relative fraction (cell type-specific/ubiquitously expressed) of gene promoters with at least the specified number of ChIA-PET interactions with other cell type-specific HOT loci. The gray shaded area represents the 95% confidence, null interval of randomly shuffled loci interactions between cell type-specific and ubiquitously expressed promoters. (F) Cumulative distribution functions indicating the cumulative fraction of loci that contain a neighboring locus with an equivalent number of DAP associations (as specified by the line color) within a given distance in base pairs.

## Notes

https://www.ncbi.nlm.nih.gov/geo/query/acc.cgi?acc=GSE104247

### Competing Interest Statement

The authors have declared no competing interest.

### Summary of Updates

Revisions made to text and figures.

